# The Dilute domain of Canoe is not essential for Canoe’s role in linking adherens junctions to the cytoskeleton but contributes to robustness of morphogenesis

**DOI:** 10.1101/2023.10.18.562854

**Authors:** Emily D. McParland, T. Amber Butcher, Noah J. Gurley, Ruth I. Johnson, Kevin C. Slep, Mark Peifer

## Abstract

Robust linkage between cell-cell adherens junctions and the actomyosin cytoskeleton allows cells to change shape and move during morphogenesis without tearing tissues apart. The multidomain protein *Drosophila* Canoe and its mammalian homolog Afadin are critical for this linkage, and in their absence many events of morphogenesis fail. To define underlying mechanisms, we are taking Canoe apart, using *Drosophila* as our model. Canoe and Afadin share five folded protein domains, followed by a large intrinsically disordered region. The largest of these folded domains is the Dilute domain, which is found in Canoe/Afadin, their paralogs, and members of the MyosinV family. To define the roles of Canoe’s Dilute domain we have combined biochemical, genetic and cell biological assays. Use of the AlphaFold tools revealed the predicted structure of the Canoe/Afadin Dilute domain, providing similarities and contrasts with that of MyosinV. Our biochemical data suggest one potential shared function: the ability to dimerize. We next generated *Drosophila* mutants with the Dilute domain cleanly deleted. Surprisingly, these mutants are viable and fertile, and CanoeΔDIL protein localizes to adherens junctions and is enriched at junctions under tension. However, when we reduce the dose of CanoeΔDIL protein in a sensitized assay, it becomes clear it does not provide full wildtype function. Further, canoeΔ*DIL* mutants have defects in pupal eye development, another process that requires orchestrated cell rearrangements. Together, these data reveal the robustness in AJ-cytoskeletal connections during multiple embryonic and postembryonic events, and the power of natural selection to maintain protein structure even in robust systems.

## Introduction

Building the architecture of tissues and organs requires individual cells to work together, changing shape and moving in a coordinated way. Cell shape change requires force to be exerted on the plasma membrane. This occurs at cell-cell adherens junctions (AJs) and cell-extracellular matrix junctions. At these junctions transmembrane cadherins or integrins connect cells to one another or to the extracellular matrix, respectively. Their cytoplasmic domains then organize linker proteins that connect to the actin cytoskeleton on which myosin motor proteins walk, generating the force required to drive shape change.

Scientists studying integrin-based adhesions have long appreciated the complexity of the mechanosensitive protein network that links integrin cytoplasmic tails to the actomyosin cytoskeleton, with dozens of components arranged in distinct layers (Case and Waterman, 2015). Our view of the linkage of cadherins to the cytoskeleton started much more simply, with a picture of a linear and direct linkage. In this picture beta-catenin bound both the cadherin tail and alpha-catenin, while alpha-catenin bound F-actin. Subsequent work revealed that this picture is significantly over-simplified (reviewed in (Perez-Vale and Peifer, 2020; Yap, Duszyc and Viasnoff, 2018). We now recognize that a much larger network of proteins mediates this linkage, including Afadin/Canoe (Cno), ZO-1/Polychaetoid, Ajuba, Vinculin, Sidekick and likely others. This network provides redundancy and thus robustness, with individual protein interactions and even some individual proteins dispensable for baseline function. The protein network is also mechanically sensitive, with the linkage strengthened by pulling force on the AJ.

Part of the robustness of the network is conferred by the fact that many proteins in it are large multidomain proteins that can bind many partners. We use Cno protein and its mammalian homolog Afadin as a model for exploring how this complex protein structure confers function. Cno and Afadin share five predicted folded protein domains, followed by a long intrinsically disordered region (Fig. 1A; (Gurley et al., 2023). At their amino terminus are two Ras-association (RA) domains, which bind to the small GTPase Rap1 (Boettner et al., 2000). Rap1 “activates” Cno by mechanisms we are only starting to understand (Boettner et al., 2003; Bonello et al., 2018; Perez-Vale et al., 2023). The central PDZ domain binds several partners, including the transmembrane junctional proteins E-cadherin (Ecad) and Nectins/Echinoid (Sawyer et al., 2009; Takahashi et al., 1999; Wei et al., 2005). Between the RA and PDZ domains are two other domains—the Forkhead-associated (FHA) and Dilute (DIL) domains--which can be recognized by sequence, but the biochemical function of which remains unclear. The C-terminal intrinsically disordered region carries one or more F-actin binding sites, the most C-terminal of which is referred to as the F-actin binding (FAB) region (Mandai et al., 1997; Sakakibara et al., 2018).

**Figure 1.**
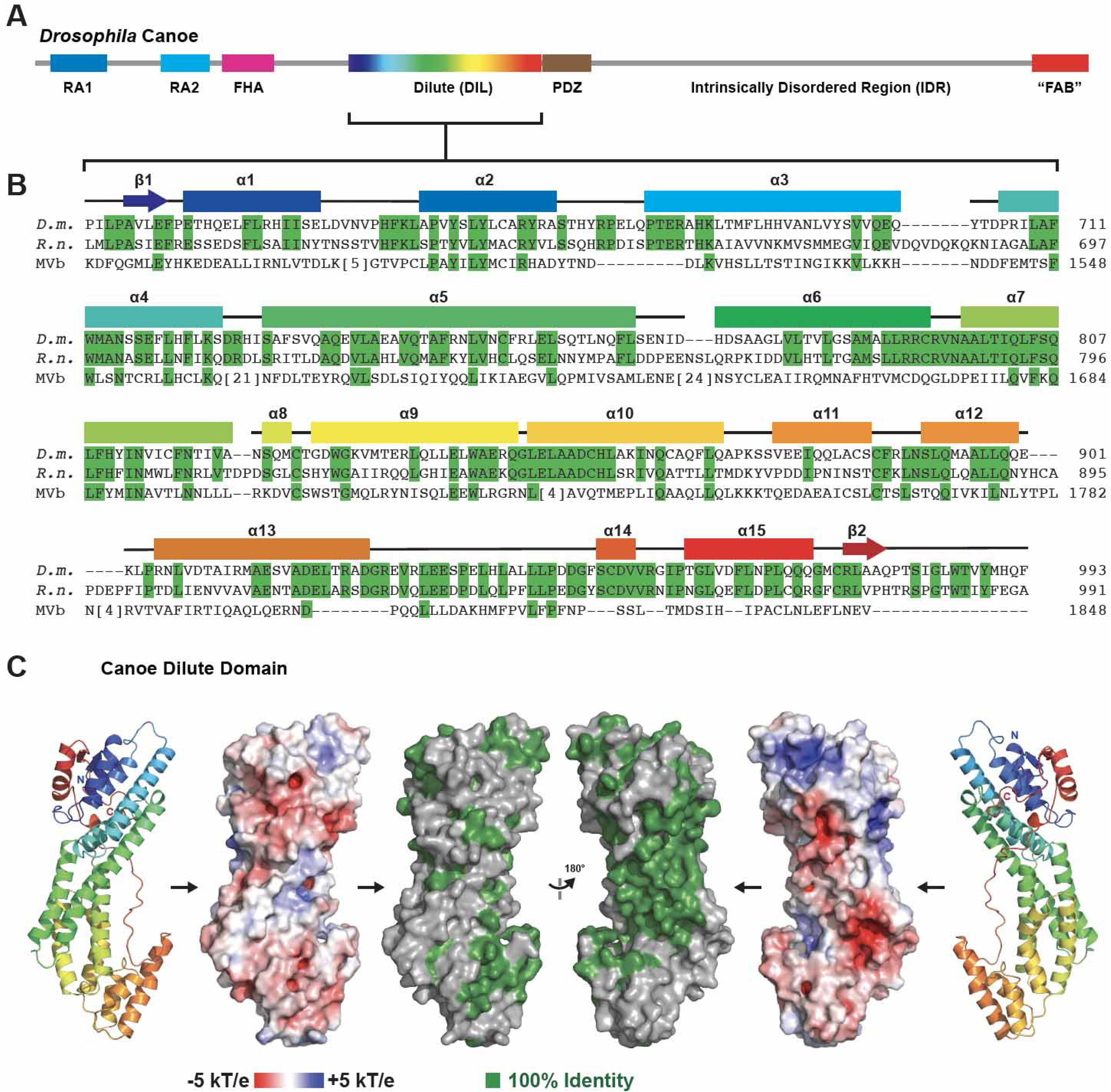
The Cno/Afadin family DIL domain has a conserved central groove. (A) Domain architecture of Drosophila Cno. The DIL domain is colored in a blue to red spectrum, representative of the secondary structure color codes in B and C. (B) Sequence alignment of Drosophila melanogaster (D.m.) Cno, Rattus norvegicus (R.n.) Afadin, and the sequence of the human MyoVb (MVb) DIL domain. Residues identical between Cno and Afadin are highlighted in green, as are the corresponding residues in MyoVb for which the identity extends. Predicted secondary structure elements of the Cno AlphaFold prediction (Varadi et al., 2022){Jumper, 2021 #5829 are shown above the alignment. (C) AlphaFold structure prediction of the Cno DIL domain shown (from left to right) in cartoon format, space-filling format showing electrostatics, and conservation (residues identical between rat Afadin and fly Cno as highlighted in B), followed by views of each after a 180° rotation about the y-axis.

We are using Drosophila’s powerful genetic tools to take apart this complex multidomain machine and analyze the function of different domains. Cno is important for almost every cell shape change or morphogenetic movement in the embryo, ranging from initial positioning of AJs (Choi et al., 2013) to apical constriction of mesodermal cells (Sawyer et al., 2009) to convergent elongation of the body axis (Sawyer et al., 2011; Yu and Zallen, 2020) to the collective cell migrations of dorsal closure and head involution (Boettner et al., 2003; Choi et al., 2011). Cno is required to strengthen junction-cytoskeletal connections at AJs under elevated force, and in its absence cytoskeletal-AJ connections are disrupted. We generated a CRISPR-based tool that allows us to replace the endogenous *cno* coding sequence with GFP-tagged mutant proteins (Perez-Vale et al., 2021). Deleting both N-terminal RA-domains nearly eliminated Cno function (Perez-Vale et al., 2021). CnoΔRA localization to forming AJs was altered, though this was restored later. However, the tension-sensitive recruitment of Cno to AJs under tension requires the RA domains. We also explored the roles of the PDZ domain and FAB region (Perez-Vale et al., 2021), hypothesizing they would be essential for function by linking Ecad to actin. However, to our surprise, both were dispensable for viability. Sensitized assays revealed that these domains reinforce AJs under tension. Together, these data and those from others support the idea that a robust AJ-cytoskeletal linkage is conferred by multivalent interactions.

Here, we focus on the DIL domain of Cno, asking what role(s) it plays in Cno function. The DIL domain was first identified in the unconventional myosin, Myosin V (MyoV), which transports vesicles (reviewed in (Wong and Weisman, 2021). MyoV’s DIL domain (also known as the Cargo-binding Domain or Globular Tail Domain) provides the cargo binding site. It has an elongated structure in which 15 amphipathic α-helices are connected by short and long loops (Fig. 1B). The MyoV DIL domain binds many partners and has multiple functions. It binds the MyoV motor domain to act as an auto-inhibitor of motor activity. It is also the docking site for multiple cargo-specific adapter proteins, including Melanophilin, Spire2, Molecules Interacting with CasL (MICAL1) and Rab interacting lysosomal protein-like 2 (RIPL2). These proteins dock on three spatially distinct regions on the DIL domain (e.g. (Wei et al., 2013)). Small Rab family GTPases also dock on the DIL domain (Pylypenko et al., 2013). Structural studies suggest that some adapters may indirectly dimerize MyoV via their DIL domain interaction (Wei et al., 2013). At least a subset of MyoV DIL domains can also dimerize directly, with two different potential modes of interaction identified (e.g. (Nascimento et al., 2013; Zhang, Yao and Li, 2016). Thus, in the protein family where it has been most closely examined the DIL domain mediates many functions, serving as a hub for both intra- and intermolecular interactions.

The only other proteins known to have DIL domains are the Cno/Afadin family, found in all animals, and their distantly related and less studied vertebrate paralogs RADIL and Rasip. The DIL domains of human Afadin and Drosophila Cno share 48% sequence identity (Gurley et al., 2023). Conservation in the vertebrate lineage is even stronger, with the DIL domains of human and zebrafish Afadin sharing 89% identity. Even the distant animal relative Trichoplax contains a Cno/Afadin relative, with a DIL domain 38% identical to that in Drosophila. Despite this strong conservation, little is known about the molecular or biological functions of the DIL domain in either mammals or Drosophila. Yeast two-hybrid protein interaction screens identified the coiled-coil protein Afadin DIL domain-Interacting Protein (ADIP) as a binding partner of Afadin that can interact with its DIL domain (Asada et al., 2003). ADIP can regulate cell migration of cultured cells, by regulating the small GTPase Rac (Fukumoto et al., 2011). Only two tests of Afadin DIL domain function have been reported, both in cell culture. In migrating mammalian cells, an Afadin mutant lacking the DIL domain did not fully support cell migration in a cultured cell wounding assay, matching the effect of ADIP knockdown (Fukumoto et al., 2011). In cultured MDCK cells manipulated to elevate junctional tension, an Afadin mutant lacking the DIL domain provided strong rescue of the gaps in cell junctions under tension caused by Afadin knockdown but did not fully rescue cell shape (Choi et al., 2016). However, this latter cell culture assay used Afadin knockdown, not knockout, and thus in this system even an Afadin mutant lacking the Rap1-binding RA domains, which provides only minimal function in Drosophila, provided significant rescue (Choi et al., 2016). To fully explore the biochemical and biological function of the DIL domain in Cno’s diverse roles, we combined biochemical, genetic and cell biological assays.

## Results

### The DIL domains of Cno/Afadin have a unique, conserved central groove

The DIL domains of Cno and Afadin are well conserved in sequence and length, with 47% of the 374 fly amino acids identical to the corresponding rat 393 amino acids (Gurley et al., 2023); Fig. 1B). The structure of the DIL domain has been solved from multiple MyoV family members, both alone and in complex with binding partners, but the structure of Cno/Afadin DIL domains has not been experimentally determined. New tools made available by the development of AlphaFold allowed us to examine the predicted structures of the DIL domain in Cno and Afadin (Fig. 1B,C, 2; (Jumper et al., 2021; Varadi et al., 2022). Cno’s DIL domain is predicted to contain 15 α-helices that, like the MyoV DIL domain, form an elongated domain. A C-terminal tail is folded back along the domain, positioning the C-terminal β2 strand anti-parallel to the N-terminal β1 strand. The region containing the N- and C-termini, which are proximal to one another, is predominantly basic, as compared to the rest of the domain, which is primarily acidic (Fig. 1C). The PDZ domain, which is just C-terminal to the DIL domain, binds membrane-associated proteins. Thus, the basic nature of the DIL domain proximal to the PDZ domain may serve to complement the negatively-charged plasma membrane.

Aligning the predicted structures of Cno and Afadin with the solved structure of MyoVb (Nascimento et al., 2013) highlights unique features of the Cno/Afadin family as well as regions where either Cno or Afadin is more like MyoVb while the other deviates (Fig. 2). One area involves the terminal regions of the domain (Fig. 2A). While Cno and Afadin are predicted to have similar structure over this region (Fig. 2A left panel), MyoVb lacks the α15 helix (Fig. 2A, left panel, red arrow), and has a larger insertion between α5 and α6 which extends away from the helices and occupies the site corresponding to the Cno/Afadin C-terminal tail (Fig. 2A right panel, lower red arrow). In Cno/Afadin, the corresponding loop occurs between helices α4 and α5, and is minimal in length. The structural alignment also highlights loops of different lengths and positioning, including 1) the Cno α2-α3 loop, which is similar to Afadin (Fig. 2A, left panel), but is extended relative to MyoVb (Fig. 2A, right panel, upper red arrow), 2) the Cno α3-α4 loop (Fig. 2B, red arrow) which is similar to Myo5b, but distinct from that of Afadin, 3) the Cno α5-α6 loop, which is extended in Afadin (Fig. 2C, red arrow), and disordered in MyoVb (Fig. 2C, the MyoV phospho loop, red circles and red dotted line), suggesting high structural plasticity (Fig. 2C), and 4) the Cno α12-α13 loop, which is similar between Afadin and MyoVb, but is shorter and distinctive in Cno (Fig. 2D, lower red arrow). The Cno α13 helix is similar to Afadin α13 as well, but the corresponding MyoVb helix is truncated from the C-terminal region (Fig. 2D, upper red arrow). Collectively, the structural comparisons highlight a common core domain with structural variations primarily focused on the length and positioning of select loops. We also examined which regions of Cno and Afadin are most highly conserved. Strong sequence identity between Cno and Afadin maps to a central groove on the domain, largely consisting of residues from α6, α7, α9, and α10 that are distinct from those in MyoV (Fig. 1B,C), suggesting a potential conserved site specific to the Cno/Afadin family for a molecular interaction.

**Figure 2.**
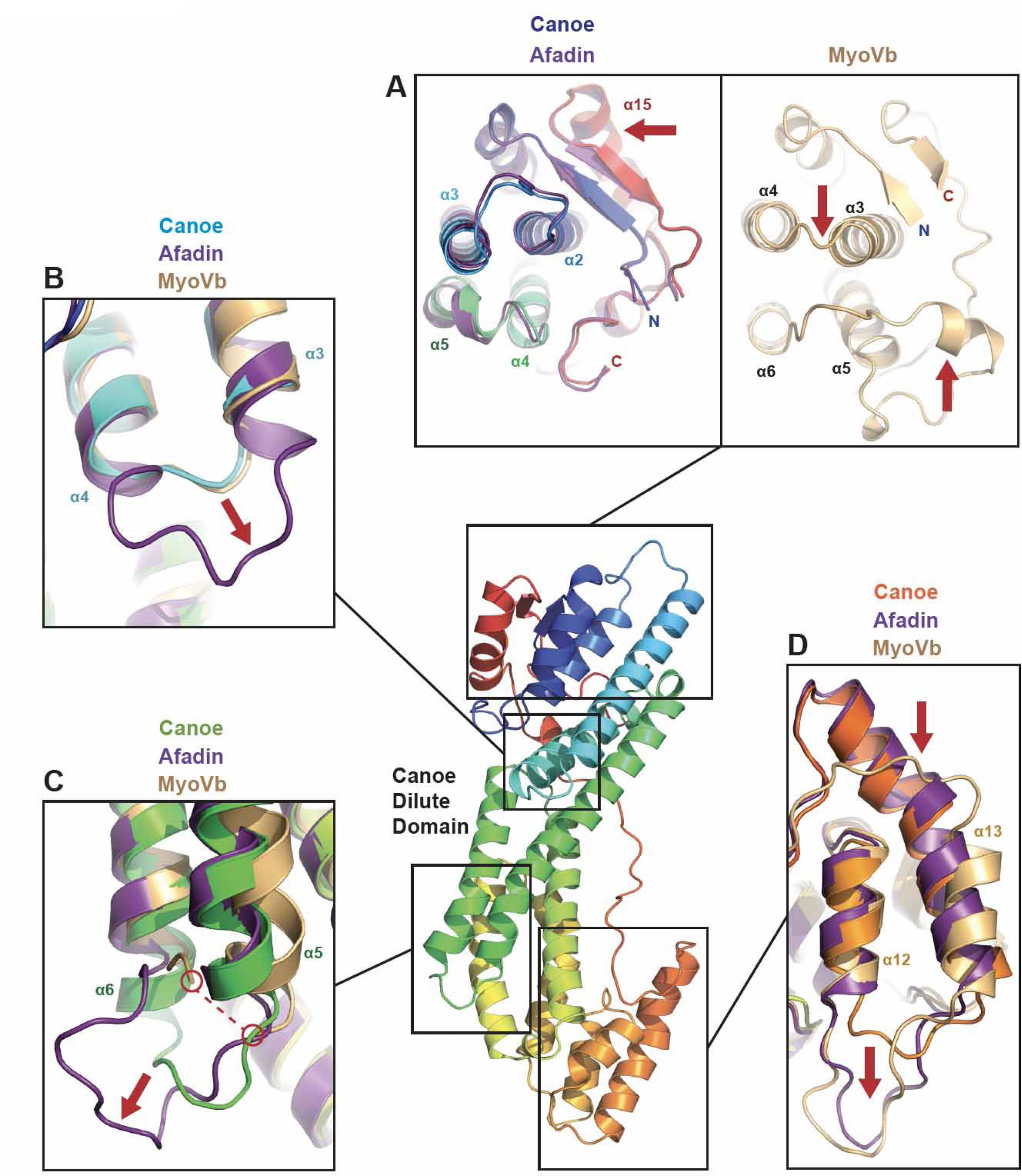
The predicted DIL domain structures of Cno and Afadin are structurally similar to the MyoVb DIL domain but have components that structurally diverge. Center: Cno DIL domain AlphaFold structure prediction, shown in cartoon format, colored as in Figure 1C. Regions boxed are shown in more detail in A-D, and include structural alignments with the predicted structure of Afadin, and the experimentally determined structure of MyoVb from PDB 4J5M (Nascimento et al., 2013)). (A) Zoom view of the top of the DIL domains (rotated 90° about the x-axis relative to the center image) after structural alignment of the Cno and Afadin (shown at left) and MyoVb (shown at right) DIL domains. Cno and Afadin have high structural homology over this region, while Myo5b has a unique helical insert (lower red arrow in right panel) between its α5 and α6 helices that occupies space which in the Cno/Afadin DIL domain structures is occupied by the domain’s C-terminal tail. Cno and Afadin have a distinct orientation of the α2-α3 loop that deviates from the corresponding MyoVb α3-α4 loop (right panel, upper red arrow). Cno and Afadin also have a C-terminal helix, α15, that is not present in the MyoVb structure (red arrow, left panel). (B) Zoom view showing structural differences in the positioning and length of the Cno and Afadin α3-α4 loop (red arrow), showing that the Cno α3-α4 loop is positioned similar to the corresponding loop in MyoVb. (C) Zoom view showing the variation in the positioning of the Cno and Afadin α5-α6 loops, which are not ordered in the MyoVb structure (MyoVb disordered loop indicated by a bridging red-dotted line). (D) Zoom view of the Cno and Afadin DIL domain α12 -α13 region, showing variation in the positioning and length of the α12-α13 loop (lower red arrow), and the different length of the α13 helix (upper red arrow) which is extended in both Cno and Afadin, but is relatively shorter in the MyoVb structure.

While the conserved central groove of the Cno/Afadin DIL domain is distinct from MyoV, we inquired whether any MyoV binding partners, for which structures were determined, occupied binding sites that overlapped the central groove (Fig. 3). We structurally aligned MyoV DIL domains in complex with a variety of binding partners (Melanophilin, Spir2, MICAL1, RILPL2, and Rab11 (Niu et al., 2020; Pylypenko et al., 2013; Pylypenko et al., 2016; Wei et al., 2013) with the predicted Cno structure (Jumper et al., 2021; Varadi et al., 2022). Modeling the sites occupied by these MyoV binding partners on the Cno DIL domain and comparing these to regions of Cno/Afadin sequence conservation revealed that none of these MyoV binding factors fully engage the conserved central groove (Fig. 3B). A region of the RILPL2 binding site partially overlaps with the conserved central groove but leaves most of the groove open (Fig. 3B, right panel). Rab11 and the peptide binding modes of Melanophilin, Spir2, and MICAL1 engage the MyoV DIL domain in regions that are not conserved between Cno and Afadin. The MyoV DIL domain is part of a large myosin motor complex, in which intramolecular interactions maintain the motor in an off state. We next examined how Cno/Afadin conservation compares to regions involved in the interdomain interactions involving MyoV’s DIL domain (Fig. 3C). In the full length MyoVa homodimer structure, the MyoVa DIL domain engages the motor domain, the coiled coil, and the sequence N-terminal to the DIL domain of the homodimeric mate. The latter interaction occurs in a mode akin to the Melanophilin, Spir2, and MICAL1 peptides (Niu et al., 2022). None of these interdomain interactions engage the conserved central groove except for the coiled coil, which has partial binding site overlap, but leaves most of the groove accessible (Fig. 3C).

**Figure 3.**
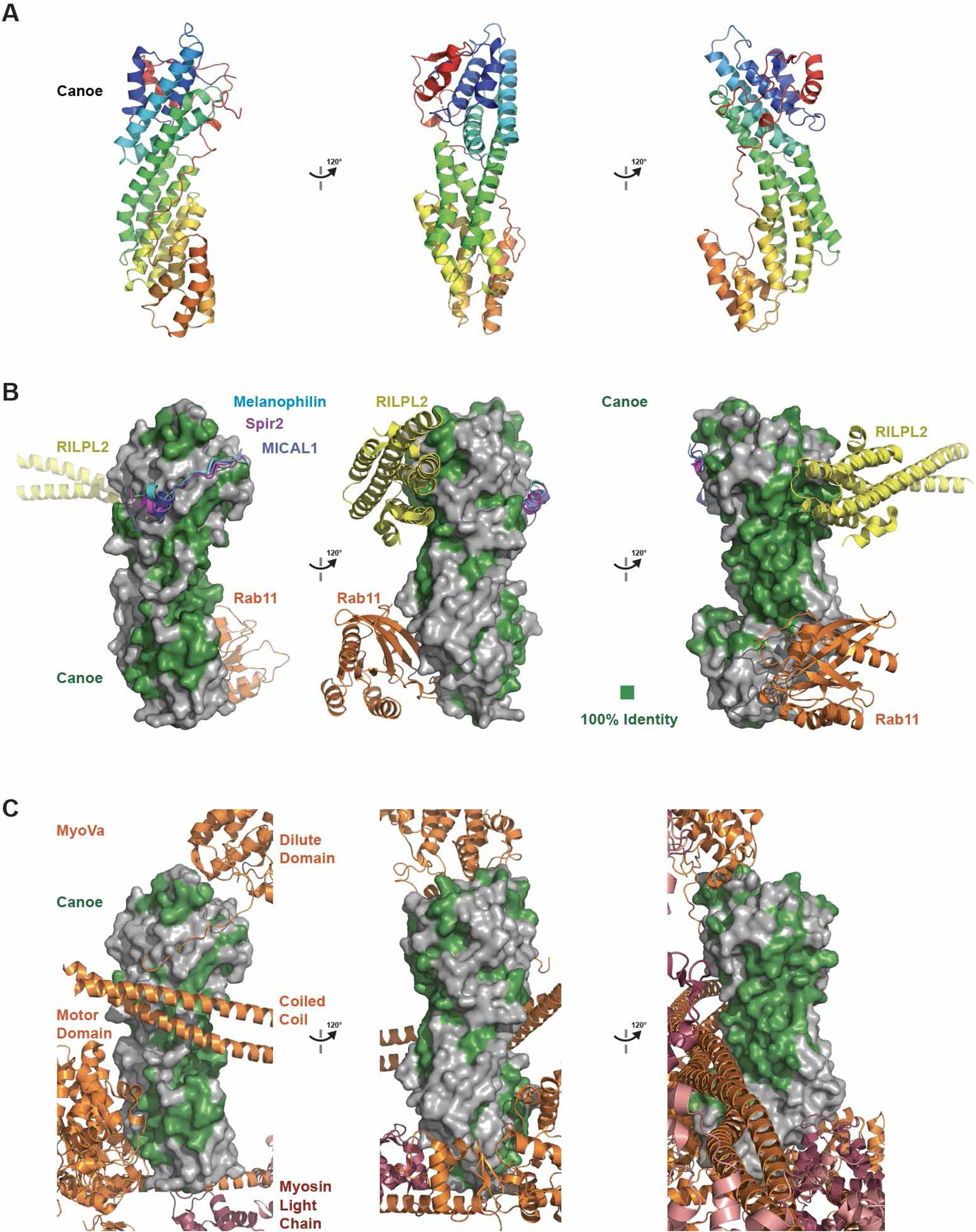
The Cno/Afadin DIL domains share a conserved central groove that is not engaged by MyoV binding factors in MyoV complex structures determined to date. (A) The AlphaFold Cno DIL domain predicted structure, shown in cartoon format in consecutive 120° rotations about the y-axis, colored as in Figure 1C. (B) Comparative analysis of where MyoV binding proteins bind the MyoV DIL domain compared with conserved residues of the Cno/Afadin family DIL domain. Cno DIL domain oriented as in A, shown in surface representation, with residues identical between D.m. Cno and R.n. Afadin colored green. MyoV binding proteins: RILPL2 (yellow, PDB 4KP3-citations for structures are in the Methods), Rab11 (orange, PDB 4LX0), Melanophilin (cyan, PDB 4KP3), Spir2 (purple, PDB 5JCY) and MICAL1 (dark blue, PDB 6KU0) are shown after structurally aligning the respective MyoV DIL domain from the complex with the Cno DIL domain. The MyoV DIL domain from each complex structure is not shown. MyoV binding proteins shown do not fully engage the DIL domain’s central groove that is highly conserved in Cno/Afadin. (C) Comparative analysis of how domains of the full length homodimeric MyoVa protein (the motor domain, coiled coil, and DIL domain) and myosin light chains engage the motor’s DIL domain compared with conserved residues of the Cno/Afadin family DIL domain. Cno DIL domain oriented as in A, shown in surface representation, with residues identical between D.m. Cno and R.n. Afadin colored green. MyoVa domains (orange) and myosin light chains (pink and red) from PDB 7YV9), are shown after structurally aligning one of the MyoVa DIL domains and the Cno DIL domain. The aligned MyoVa DIL domain is not shown, but the non-aligned MyoVa DIL domain of the homodimer is shown. MyoVa interdomain interactions that involve the DIL domain do not fully engage the DIL domain’s central groove that is highly conserved in Cno/Afadin.

### The DIL domain of Cno can dimerize in vitro

Data from both mammalian Afadin and Drosophila Cno suggest that these proteins can dimerize or oligomerize (Bonello et al., 2018; Mandai et al., 1997). The structural similarities with MyoV’s DIL domain, which can dimerize, encouraged us to explore whether the DIL domain of Cno shared the ability to dimerize. To test this, we produced the DIL domain of Cno in E. coli and purified it (Fig. 4A)]. We then used Size Exclusion Chromatography and Multi-Angle Light Scattering (SEC-MALS) to determine the molecular weight of the Cno DIL domain. Strikingly, the Cno DIL domain migrated as both a monomer and a dimer (Fig. 4B). This suggests that one function of the DIL domain may be to mediate Cno dimerization.

**Figure 4.**
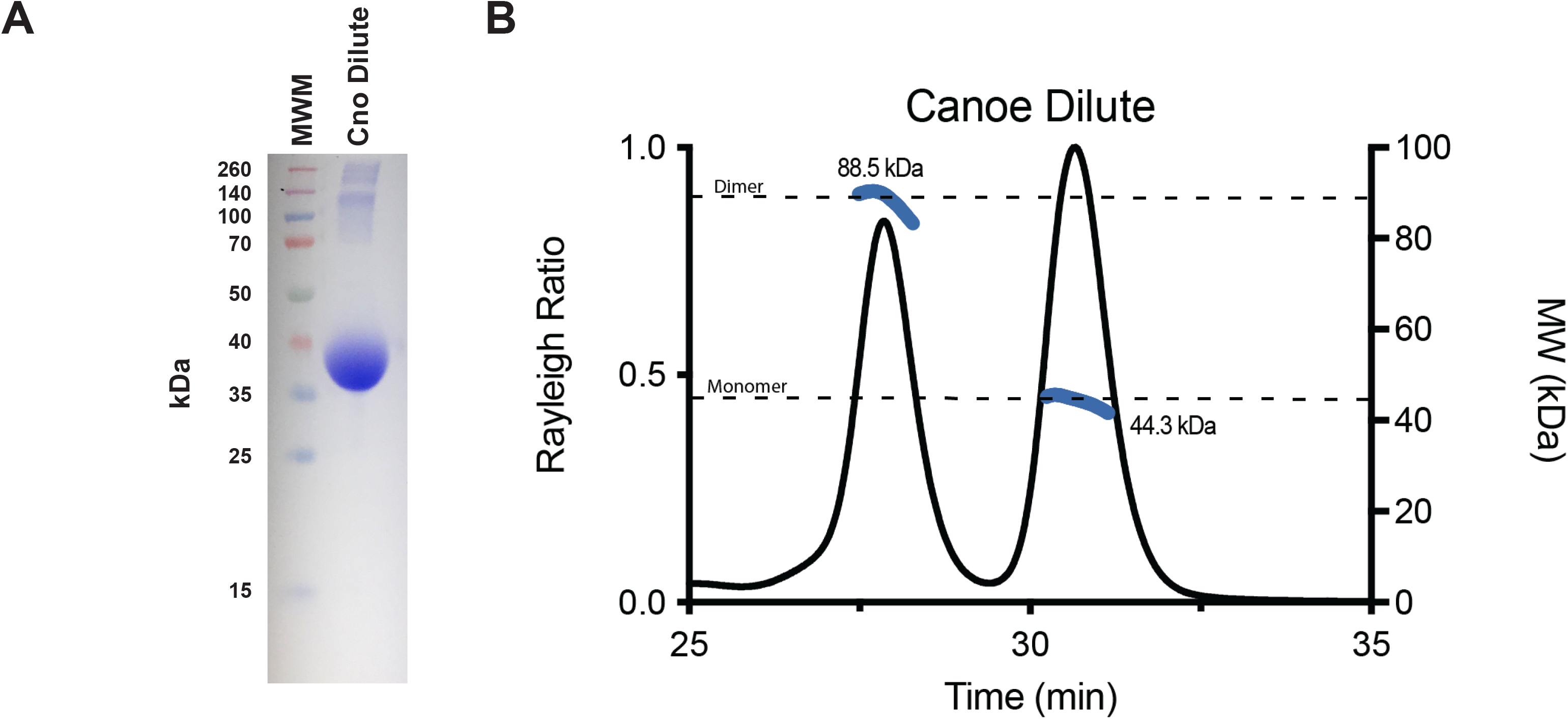
The Cno DIL domain can homodimerize in vitro. (A) SDS PAGE gel analysis of purified Cno DIL domain (aa 613-1006), FW 44.5 kDa, which runs anomalously at ∼37 kDa. A small amount of contaminants, greater than 120 kDa in size, are present. (B) SECMALS analysis of the Cno DIL domain (aa 613-1006) shows both a monomeric and a dimeric population. Rayleigh ratio (left y-axis, solid black line) and experimentally determined molecular weight (right y-axis, blue lines) are indicated relative to elution time (x-axis). The formula mass of a monomer and a homodimer are indicated with dashed black lines. 27% of the injected DIL domain eluted as a homodimer (left peak) while 73% eluted as a monomer (right peak).

### Creating a mutant to test the function of Cno’s DIL domain in Drosophila

We next turned to testing DIL domain function in vivo, by precisely deleting it from Cno protein. We designed our mutant guided by the AlphaFold predictions of domain structure (Fig. 1C), deleting amino acids 613-993 of Drosophila Cno—our deletion started in a poorly conserved region, predicted to be disordered, 11 amino acids N-terminal to the sequence we defined as the DIL domain and ended 5 amino acids before the C-terminal end of the DIL domain, to avoid inadvertent effects on the adjacent PDZ domain (Fig. 5A). We then used a system we established in Drosophila that allows us to place GFP-tagged versions of the *cno* gene, wildtype or with site-directed mutants, into a modified version of the *cno* locus, in which most of the *cno* protein coding sequence is deleted (the ΔΔ background; Fig. 5B; (Perez-Vale et al., 2021). We inserted the modified coding sequence with the DIL domain deleted in place of *cno’s* second exon, using site-specific recombination via phiC31 integrase (Fig. 5B; (Bischof et al., 2007)—the inserted sequence also carries the *white* gene as a selectable marker. Thus, inserted *cno* transgenes are expressed under endogenous gene control at the right times and places. Insertion of a GFP-tagged wildtype Cno fully rescued viability and fertility (Perez-Vale et al., 2021). We verified the accuracy of our genetic modifications with PCR amplification of the *cno*Δ*DIL* genomic locus from transgenic Drosophila, using primer pairs that distinguished the wildtype and altered gene (Fig. 5C), and sequenced the amplified product across the span of the deletion. We refer to this mutant as *cno*Δ*DIL*.

**Figure 5.**
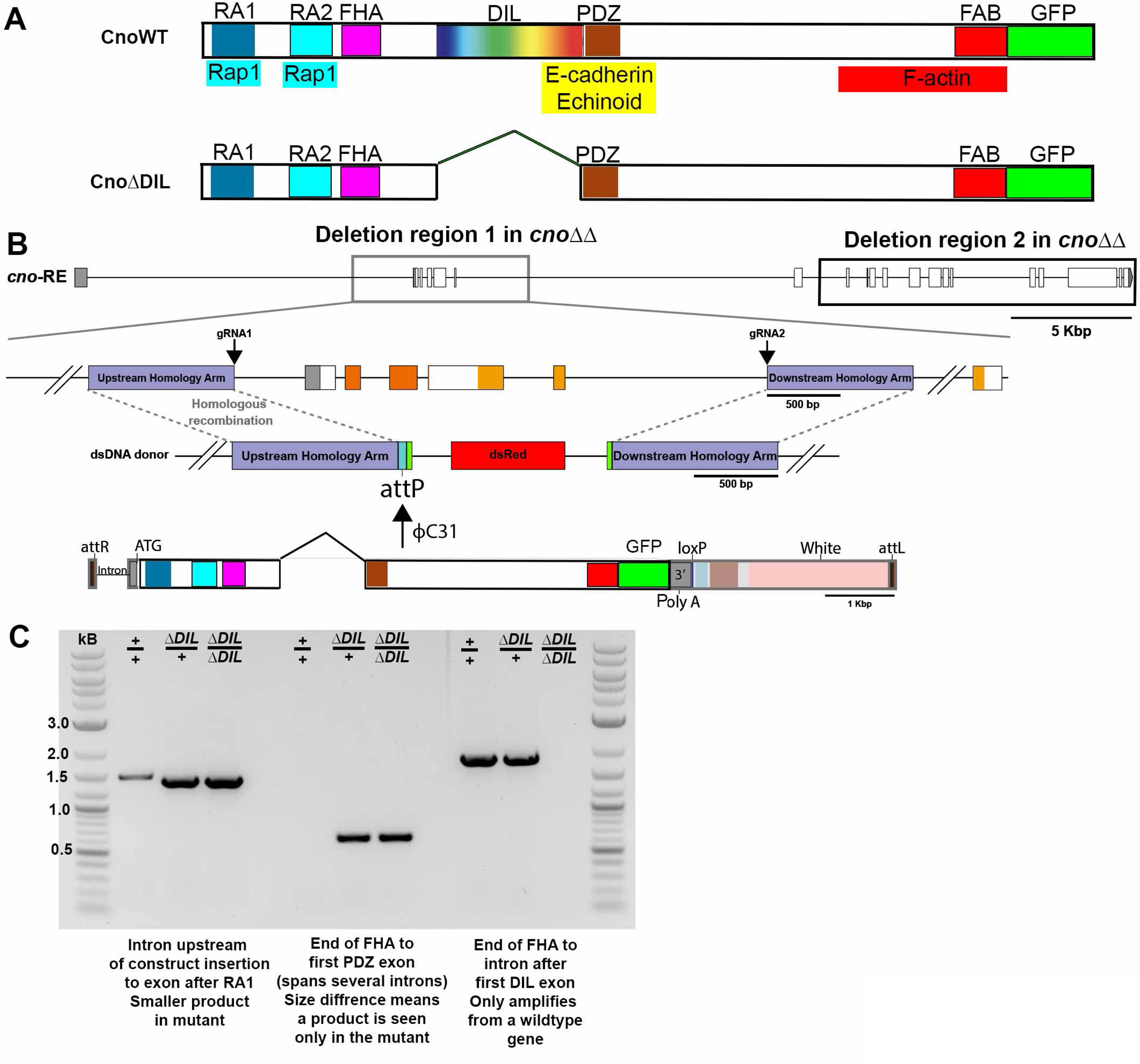
Generating a mutant to assess the function of the Cno DIL domain. (A) Diagram of the Cno protein and the CnoΔDIL mutant. (B) Strategy for generating *cno*Δ*DIL.* We started with the *cno*ΔΔ chromosome, which has most of the *cno* coding sequence deleted and has an attP site near the 3’ end of the first intron. The modified *cno* coding sequence with the DIL domain deleted and a C-terminal GFP-tag was inserted in place of the second exon of *cno* using site-specific recombination via phiC31 integrase. (C) PCR reactions confirming the correct mutation—details are in the Methods.

### CnoΔDIL protein accumulates at levels similar to wildtype Cno

Our previous site-directed mutants accumulated at levels similar to those of wildtype Cno (Perez-Vale et al., 2021). CnoΔDIL is recognized by both our standard anti-Cno antibody, which recognizes a region in the IDR (Sawyer et al., 2009), and with antibodies to the C-terminal GFP tag. We generated embryonic protein extracts from both early (1-4 hr) or mid-stage (12-15 hr) embryos and examined them by immunoblotting with anti-Cno and anti-GFP antibodies. We used antibodies to alpha-tubulin as a loading control. We examined stocks either heterozygous or homozygous for *cno*Δ*DIL,* with wildtype endogenous Cno and GFP-tagged wildtype Cno, which accumulates at wildtype levels (Perez-Vale et al., 2021) as standards. CnoΔDIL accumulates at levels similar to wildtype endogenous Cno and to GFP-tagged wildtype Cno in both early and mid-stage embryos (Fig 6A,B). Quantifying protein levels relative to GFP-tagged wildtype Cno using multiple samples verified that CnoΔDIL accumulates at essentially wildtype levels (Fig. 6C), validating its use to examine Cno DIL domain function.

**Figure 6.**
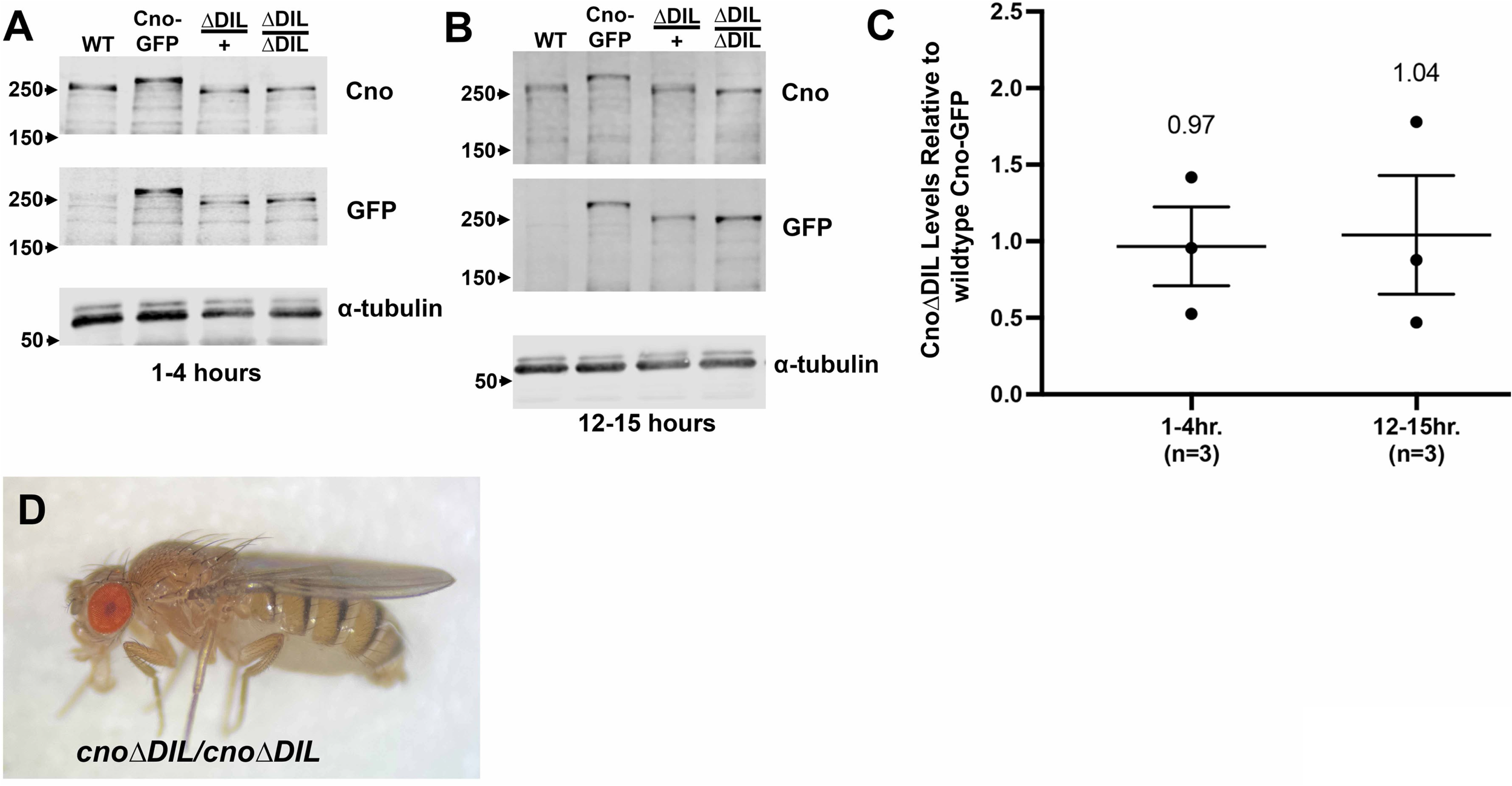
CnoΔDIL protein accumulates at normal levels and *cno*Δ*DIL* mutants are viable. (A,B) Embryonic protein extracts of the indicated ages, immunoblotted with antibodies to Cno, GFP, or alpha-tubulin as a loading control. *cnoWTGFP* embryos were a positive control for GFP antibody. Because of the deletion, GFP-tagged CnoΔDIL protein runs at a similar apparent MW to wildtype Cno. (C) Calculated levels of CnoΔDIL relative to wildtype Cno. Three replicates are shown with the broad line illustrating the mean value. D. Homozygous *cno*Δ*DIL* mutant.

### Deleting Cno’s DIL domain does not compromise viability or fertility

*cno* null mutants are zygotically embryonic lethal (Jürgens et al., 1984; Sawyer et al., 2009). The Rap1-binding RA domains are essential for viability, while the PDZ domain and the C-terminal FAB region are dispensable for viability and largely or completely dispensable for fertility (Perez-Vale et al., 2021). We thus assessed whether the conserved DIL domain is essential for viability. After obtaining *cno*Δ*DIL* flies, we outcrossed them to a wildtype stock (carrying the *y* and *w* mutations), selecting for the *w^+^* gene introduced into the *cno locus.* After multiple generations of outcrossing, we created a Balanced stock using a third chromosome Balancer chromosome and examined whether homozygous *cno*Δ*DIL* adults could be obtained. We obtained homozygous adults (Fig. 6D) and created a homozygous mutant stock, which we once again verified by PCR (Fig. 5C) and by Western blotting (Fig 6A,B). We saw no significant embryonic lethality of zygotic mutant embryos (2% lethality; n= 497 embryos). In our earlier analyses, we found that while *cno*Δ*FAB* are viable and fertile, embryonic viability of maternal/zygotic mutants is reduced (Perez-Vale et al., 2021). We thus also assessed whether this was true for *cno*Δ*DIL.* We also saw no significant embryonic lethality of maternal/zygotic mutants (6%; n=199). Thus, deleting Cno’s DIL domain does not compromise viability or fertility.

### CnoΔDIL protein localizes to and supports initial assembly of AJs

We next assessed the role of the DIL domain in Cno protein localization, and looked closely at developing embryos to determine if there were defects in cell junction assembly and maintenance, or cell shape changes subtle enough to be compatible with viability. Cno localizes to embryonic cell-cell AJs from their initial assembly (Bonello et al., 2018; Sawyer et al., 2009). We visualized CnoΔDIL using either the GFP tag or, in homozygous or hemizygous mutants, antibodies to Cno (images shown include either embryos maternally and zygotically mutant for *cno*Δ*DIL* or embryos transheterozygous for *cno*Δ*DIL* and our protein null allele *cno^R2^*).

We first examined Cno localization and function as AJs assemble during cellularization. The core AJ complex, including Armadillo (Arm=beta-catenin), localizes to spot AJs that assemble near the apical end of the invaginating membrane and are positioned around the apical perimeter of forming cells (Harris and Peifer, 2004); Fig.7A’, B’ green arrow), with some modest enrichment in tricellular junctions (TCJs; Fig. 7A, A’ green arrows). The cadherin-catenin complex also localizes along the lateral cell membrane and is enriched in basal junctions at the tip of the invaginating membrane (Fig. 7B’ red arrow). Both wildtype Cno and Bazooka (Baz=fly Par3) localize with Arm in forming apical junctions (Fig.7A”, B” green arrows; Fig. 7C, C’ green arrows), but are absent from basal junctions Fig.7A”, B” red arrows; Fig. 7C, C’ red arrows). At this stage Cno is significantly enriched in TCJs relative to the bicellular spot AJs (Fig. 7A,A” green vs red arrows;(Bonello et al., 2018). During cellularization Cno is required for correct initial positioning of AJs—in its absence both Arm and Baz are no longer enriched in nascent junctions and instead localize all along the lateral membrane (Choi et al., 2013).

**Figure 7.**
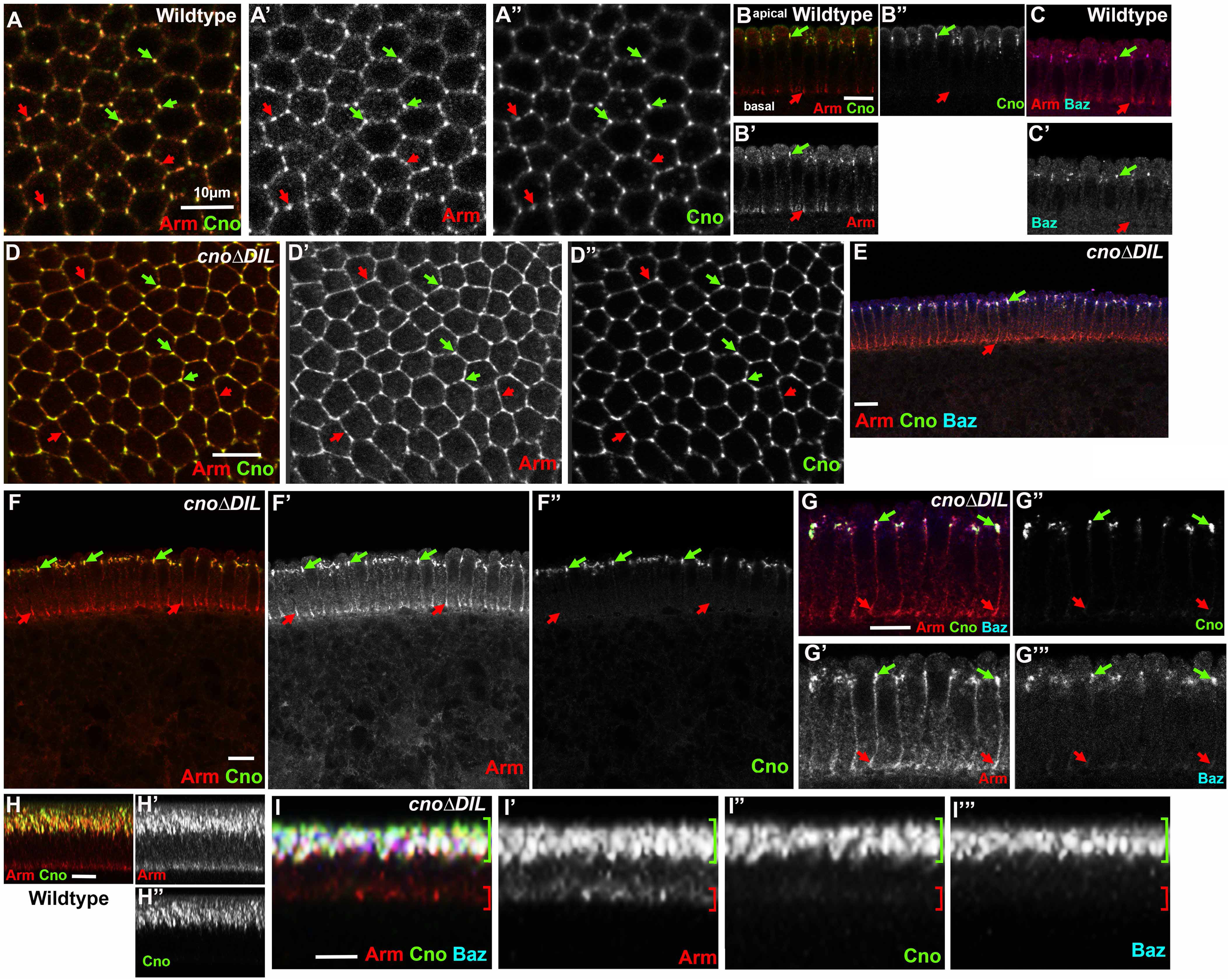
CnoΔDIL localizes and functions correctly as AJs assemble during cellularization. Late cellularization embryos, antigens and genotypes indicated. (A,D). Apical junction views of wildtype (A) or maternal/zygotic *cno*Δ*DIL* mutant embryos (D). In wildtype Arm localizes to all spot AJs (A, red arrows), with some enrichment at TCJs (A, green arrows) while Cno is substantially enriched at TCJs. CnoΔDIL protein localizes normally, with strong enrichment at TCJs (D, green vs red arrows). (B,C,E-G) Cross sections, apical up. (B,C) In wildtype Arm localizes to both nascent apical junctions (green arrows) and basal junctions (red arrows), while Cno and Baz are restricted to apical junctions. (E-G) In maternal/zygotic *cno*Δ*DIL* mutant embryos Arm and Baz localize normally, and CnoΔDIL is properly localized to apical junctions. (H-I) Maximum-intensity projections of multiple cross sections. This imaging method emphasizes the enrichment of Arm, Cno, and Baz to apical junctions in wildtype (H) and confirms that CnoΔDIL retains normal localization and function during cellularization (I).

We thus examined CnoΔDIL localization and function during cellularization. Like wildtype Cno, CnoΔDIL localized to spot AJs but was more substantially enriched at TCJs relative to Arm (Fig. 7D,D” red vs green arrows). Along the Z-axis, CnoΔDIL remained localized along with Arm to nascent apical junctions (Fig. 7E, F,G green arrows) as membranes invaginated, while Arm also localized to the basal junctions at the leading edge of the invaginating membrane (Fig. 7E,F, G, red arrows). Baz also remained enriched in nascent apical junctions in *cno*Δ*DIL* mutants (Fig. 7D’”). Correct apical enrichment of all three proteins in wildtype is emphasized in maximum X/Z intensity projections of multiple cells (Fig. 7H), and similar projections revealed parallel apical enrichment in *cno*Δ*DIL* mutants (Fig. 7E). Thus CnoΔDIL retained normal localization and function during cellularization—in these properties it resembled CnoΔFAB and CnoΔPDZ but differed from CnoΔRA (Perez-Vale et al., 2021).

### CnoΔDIL is correctly enriched at AJs under elevated tension and supports morphogenesis

As gastrulation begins, the germband elongates along the anterior-posterior (AP) axis and narrows in the dorsal-ventral (DV) axis. Consistent with the viability of *cno*Δ*DIL* mutants, germband extension proceeded without apparent defects, and like endogenous Cno, CnoΔDIL remained localized to the AJ as they matured (Fig. 8A,E). During this process, AJ and cytoskeletal proteins become planar polarized across the epithelium. Myosin and F-actin become enriched at AP cell borders, forming contractile cables that constrict those boundaries and rearrange cells (reviewed in (Perez-Vale and Peifer, 2020). Meanwhile, Baz, Ecad, Arm and Polychaetoid (Pyd=ZO-1) become enriched at DV cell borders, opposite to myosin. At stage 7 wildtype Cno is slightly enriched at AP borders relative to DV borders (Sawyer et al., 2011), similar to myosin, and this enrichment is also apparent along aligned AP borders at stage 8, as more dorsal cells round up to divide. Cno also remains enriched at TCJs during germband extension (Perez-Vale et al., 2021). This Cno enrichment at TCJs is a response to elevated tension (Yu and Zallen, 2020). We thus looked at CnoΔDIL localization, examining enrichment at TCJs and AP borders. CnoΔDIL remained clearly enriched at TCJs (Fig. 8B, arrows); quantification revealed its enrichment there was similar to that of wildtype Cno (Fig. 8K). Similarly, CnoΔDIL retained a slight enrichment at AP borders at stage 7 (Fig 8C, green vs red arrows) and along aligned AP borders at stage 8 (Fig. 8F, green arrows). Quantification confirmed this subtle enrichment at aligned AP borders (Fig. 8L). Once again, these properties of CnoΔDIL were similar to those of CnoΔFAB and CnoΔPDZ but different from those of CnoΔRA, which lost enrichment at TCJs and reversed enrichment to become elevated at DV rather than AP borders (Perez-Vale et al., 2021). Cno function is also critical to restrain Baz planar polarity. In its absence Baz is virtually lost at AP borders and thus becomes highly enriched at DV borders, and often is restricted to the central region of those borders (Sawyer et al., 2011). In contrast, in *cno*Δ*DIL* mutants Baz localized all around cells at stages 7 and 8, with moderate enrichment on DV borders (Fig 8D; red versus green arrows; Fig. 8F”’).

**Figure 8.**
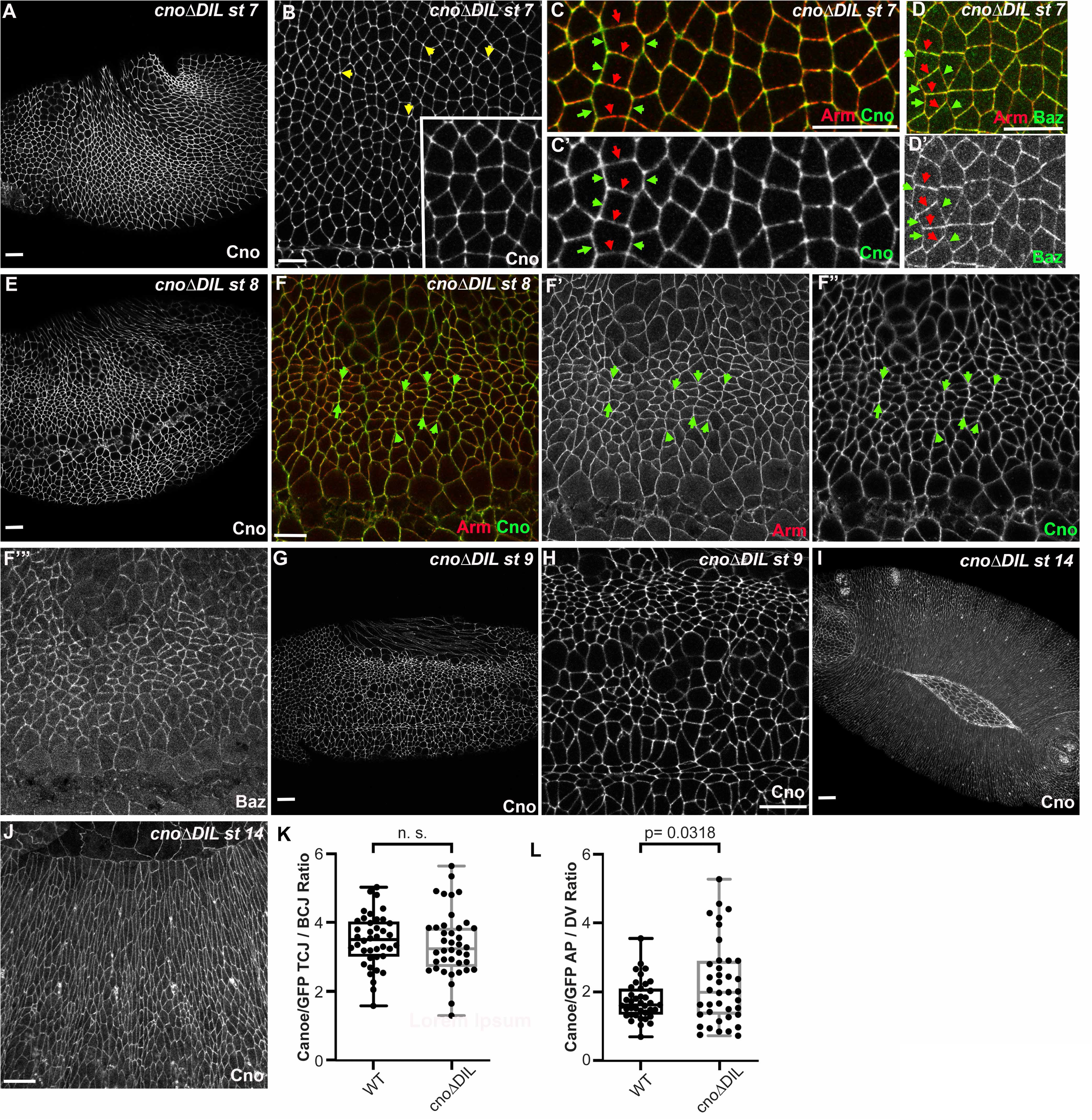
CnoΔDIL localizes and functions correctly during embryonic morphogenesis. Maternal/zygotic *cno*Δ*DIL* mutant embryos, anterior left, dorsal up, stages, genotypes and antigens indicated. (A-D) Stage 7 embryos. (A, B) CnoΔDIL localizes to AJs (A) and is enriched at TCJs (B, arrows and inset). (C) CnoΔDIL is slightly enriched at aligned AP borders (green arrows) vs DV borders (red arrows). AJ defects are not observed. (D) Baz remains localized all around the cells with some polarization to DV borders, rather than exhibiting the accentuated planar polarity seen in *cno* null mutants. (E,F) Stage 8. CnoΔDIL continues to localize to AJs (E), with some enrichment at aligned AP borders (F”, arrows). AJ defects are not observed (F’) and Baz continues to localize all around the cell (F’”). (G-J). CnoΔDIL continued to localize to AJs at stage 9 (G,H) and during dorsal closure (I,J) and no defects in morphogenesis were observed. (K) Quantification of enrichment of wildtype Cno or CnoΔDIL at TCJs. (L) Quantification of enrichment of wildtype Cno or CnoΔDIL at aligned AP vs DV cell borders.

We also looked at CnoΔDIL localization after the completion of germband extension, during stage 9 (Fig. 8G,H), and during dorsal closure (stage 14; Fig. 8I,J) and saw no differences from the localization of wildtype Cno and no deviations from normal progression of development. Taken together these data reveal that *cno*Δ*DIL* mutants retain full function in positioning nascent AJs during cellularization, in supporting cell shape changes and protein planar polarity during germband extension, and in completing dorsal closure, all events known to require Cno function. Furthermore, CnoΔDIL protein localizes to AJs and is correctly enriched at cell junctions under elevated tension.

### Deleting the DIL domain reduces Cno function when Cno protein levels are reduced

Given the strong conservation of the DIL domain in Cno/Afadin relatives in distantly related animals, we were surprised at its apparent dispensability. In our earlier analysis of *cno*Δ*PDZ* and *cno*Δ*FAB* mutants, we tested protein function in a sensitized situation in which we reduced levels of the mutant proteins. To do so, we made each mutant heterozygous with our canonical protein-null *cno* allele, *cno^R2^* (Sawyer et al., 2009). *cno^R2^*is zygotically embryonic lethal, so in a cross of *+/cno^R2^* parents, the 25% of embryos who are *cno^R2^/cno^R2^* die as embryos, while the *+/cno^R2^* and *+/+* progeny from this cross are fully viable. However, due to the strong maternal contribution of wildtype Cno, while *cno* maternal/zygotic mutants have defects in most morphogenetic movements, most *cno^R2^* homozygous zygotic mutants only have mild defects in head involution but few defects in other morphogenetic events like dorsal closure (Gurley et al., 2023; Sawyer et al., 2009). This creates a sensitized situation where small reductions in maternal Cno function can enhance these defects. Replacing wildtype Cno with either *cno*Δ*PDZ* or *cno*Δ*FAB* substantially enhanced the zygotic cuticle phenotype of *cno^R2^*homozygous zygotic mutants and led to reduced viability of *cno*Δ*PDZ/cno^R2^* or *cno*Δ*PDZ/cno^R2^* progeny (Perez-Vale et al., 2021), revealing that CnoΔFAB and CnoΔPDZ do not provide fully wildtype function. We thus implemented this sensitized genetic test to examine *cno*Δ*DIL*.

We first needed to determine if *cno*Δ*DIL/cno^R2^* transheterozygotes are adult viable. We obtained both male and female *cno*Δ*DIL/cno^R2^* transheterozygotes at Mendelian ratios (33% of progeny from a cross of *cno*Δ*DIL/TM3 x cno^R2^/TM3* (n=163 adults; Fig. 9A). We then used our sensitized test to determine whether *cno*Δ*DIL* provides wildtype Cno function. When we crossed *cno*Δ*DIL/cno^R2^*males and females, we observed slightly elevated lethality relative to the cross of *+/cno^R2^* males and females (Fig. 9B; 33%; n=1250 embryos vs 28% for *+/cno^R2^*parents; n=1482 embryos). This suggested some *cno*Δ*DIL/cno^R2^* progeny were dying in our experimental cross. However, this was not as elevated as we had observed with *cno*Δ*PDZ* or *cno*Δ*FAB* crosses, in which most of the *cno*Δ*PDZ/cno^R2^* progeny and all of the *cno*Δ*PDZ/cno^R2^* progeny died as embryos (Perez-Vale et al., 2021).

**Figure 9.**
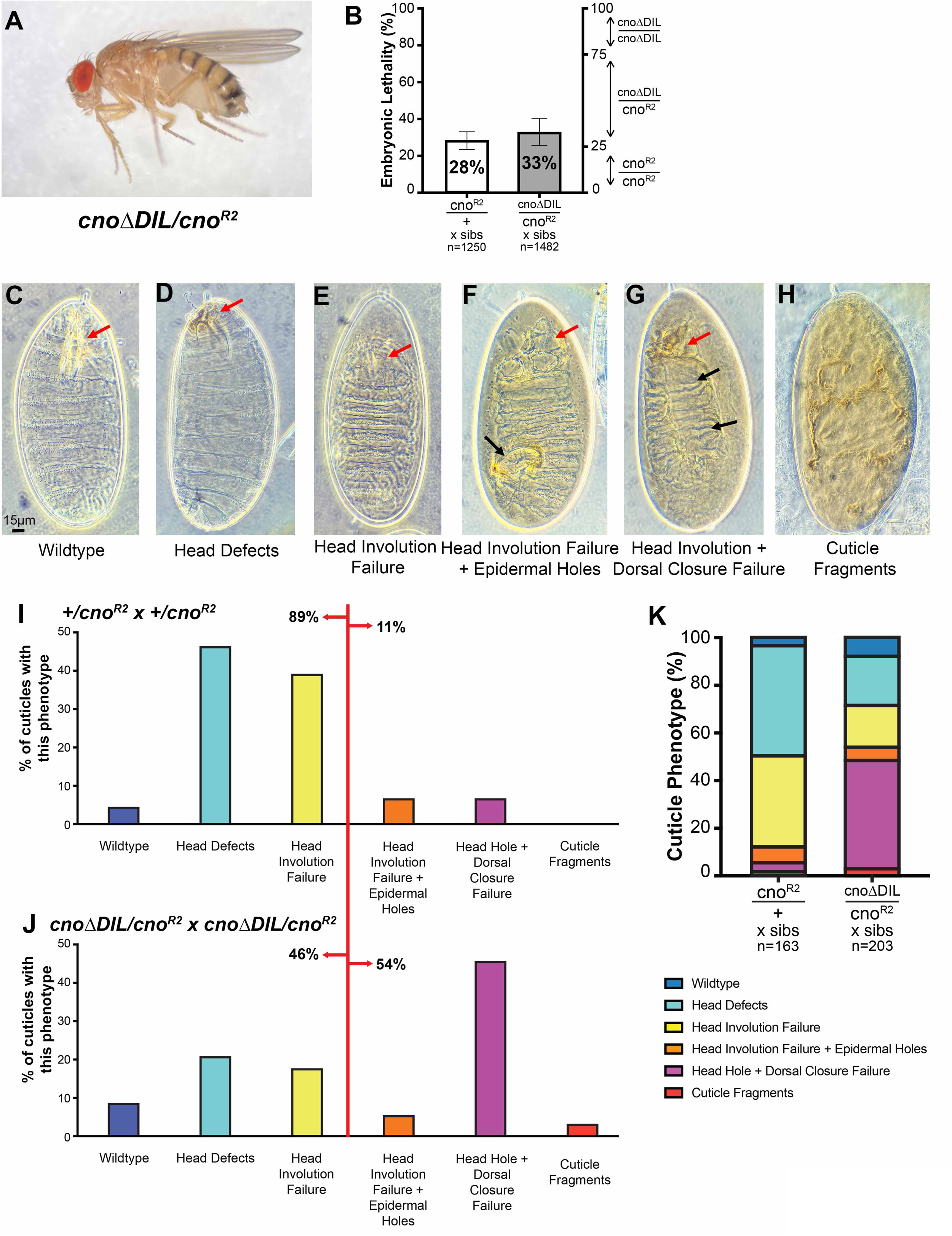
CnoΔDIL is not required for viability, but a sensitized assay reveals roles in morphogenesis. (A) *cno*Δ*DIL/cno^R2^*mutant. (B) *cno^R2^* is zygotically embryonic lethal, so 25% of progeny of heterozygous parents are expected to die as embryos. We observed 28% lethality. Embryonic lethality of the progeny of *cno*Δ*DIL/cno^R2^*parents was slightly elevated (33%), suggesting some *cno*Δ*DIL/cno^R2^* embryos die. C-H. Potential cuticle phenotypes of *cno* mutants, becoming more severe from left to right. Red arrows = head defects, black arrows = epidermal defects. (C) Wildtype. Dorsal closure and head involution are completed correctly and the head skeleton is formed correctly (arrow). (D) Defects in the head skeleton (arrow). (E) Head involution failed, leading to a hole in the anterior end of the cuticle (arrow). (F) Head involution failed red arrow) and there were holes in the dorsal or ventral cuticle (black arrow). (G) Head involution (red arrow) and dorsal closure failed, leaving the dorsal cuticle open (black arrows). (H) More severe defects in epidermal integrity. (I, K left) In the *+/cno^R2^* cross, most progeny only have defects in head involution. (J, K right) In the *cno*Δ*DIL/cno^R2^* cross, almost half of the progeny exhibit failure of both head involution and dorsal closure.

We next examined whether having a maternal contribution of CnoΔDIL protein rather than wildtype Cno would enhance the embryonic morphogenesis phenotypes of the *cno^R2^* homozygous zygotic mutant progeny. We first assessed the larval cuticle, which provides a sensitive readout of many aspects of embryonic morphogenesis requiring cell adhesion and the connection to the cytoskeleton, ranging from germband extension and retraction to dorsal closure to head involution. The cuticle of wildtype embryos is intact, has a well-developed head skeleton, the result of successful head involution, and is closed dorsally (Fig. 8C). In most *cno^R2^* zygotic mutants derived from *+/cno^R2^* parents, germband extension, retraction and dorsal closure go to completion, such that the only defects in most embryos are in head involution, leading to a disrupted head skeleton (Fig. 9E, F; (Gurley et al., 2023)—89% of embryos were in these categories, while only 11% had more severe phenotypes (Fig. 9F-H; quantified in 9I,K). In contrast, in crosses of *cno*Δ*DIL/cno^R2^* parents, many more embryos exhibited defects in or complete failure of dorsal closure (Fig 9F,G), with 54% of the embryos in more severe categories (quantified in Fig 9I,K). This enhancement of phenotypic severity strongly suggests that CnoΔDIL does not provide fully wildtype Cno function.

We also stained embryos to directly observe cell shape changes and morphogenetic movements. The strong maternal contribution of Cno only begins to diminish during dorsal closure, and thus embryos zygotically homozygous mutant for *cno^R2^* from heterozygous wildtype mothers only begin to exhibit morphogenetic defects at that stage (Choi et al., 2011; Gurley et al., 2023; Sawyer et al., 2009). Most *cno^R2^* zygotic mutant embryos complete dorsal closure and only exhibit defects in head involution (Gurley et al., 2023). Our cuticle data revealed that when the protein contributed maternally was CnoΔDIL, the *cno^R2^*zygotic mutant cuticle phenotype was enhanced, with many embryos exhibiting defects in dorsal closure (Fig. 9). In wildtype embryos, dorsal closure is driven by amnioserosal apical constriction, leading edge cable contraction and leading-edge zipping at the canthi. Together these ensure that closure is complete before amnioserosal apoptosis (Fig. 10A, B). In contrast, in many embryonic progeny of crosses between *cno*Δ*DIL/cno^R2^* parents dorsal closure clearly failed, with separation of the amnioserosa and leading edge (Fig. 10C, red arrows) or complete failure to close before amnioserosal apoptosis (Fig. 10D, red arrows)—this was consistent with our cuticle data. We also observed other defects we previously observed in *cno* mutants (Manning et al., 2019), such as persistent deep segmental grooves (Fig. 10C, D yellow arrows) and uneven cell shapes at the leading edge during dorsal closure (Fig. 10E vs F), with some cells hyperconstricted and others splayed open (Fig. 10F, red vs yellow arrows). In some embryos from the *cno*Δ*DIL/cno^R2^* cross we observed one additional earlier defect which was not previously seen in *cno* zygotic mutants but is present in maternal/zygotic *cno* mutants: defects in completion of mesoderm invagination. We observed these defects in 36% of embryos from the *cno*Δ*DIL/cno^R2^*cross. Defects included both mild defects along the ventral midline in which some mesoderm was exposed (Fig. 10G vs H; 9/39 embryos scored) and more severe defects in which the ventral furrow remained open at the anterior or posterior end (Fig. 10I vs J, K); 5/39 embryos). The frequency of defects was substantially higher than what we observed in the *+/cno^R2^* cross (6%= 3/51 had mild defects, 0/51 were anterior open) or in wildtype embryos (6%; 2/34 had mild defects, 0/34 were anterior open). Taken together, these data reveal that CnoΔDIL does not retain fully wildtype function.

**Figure 10.**
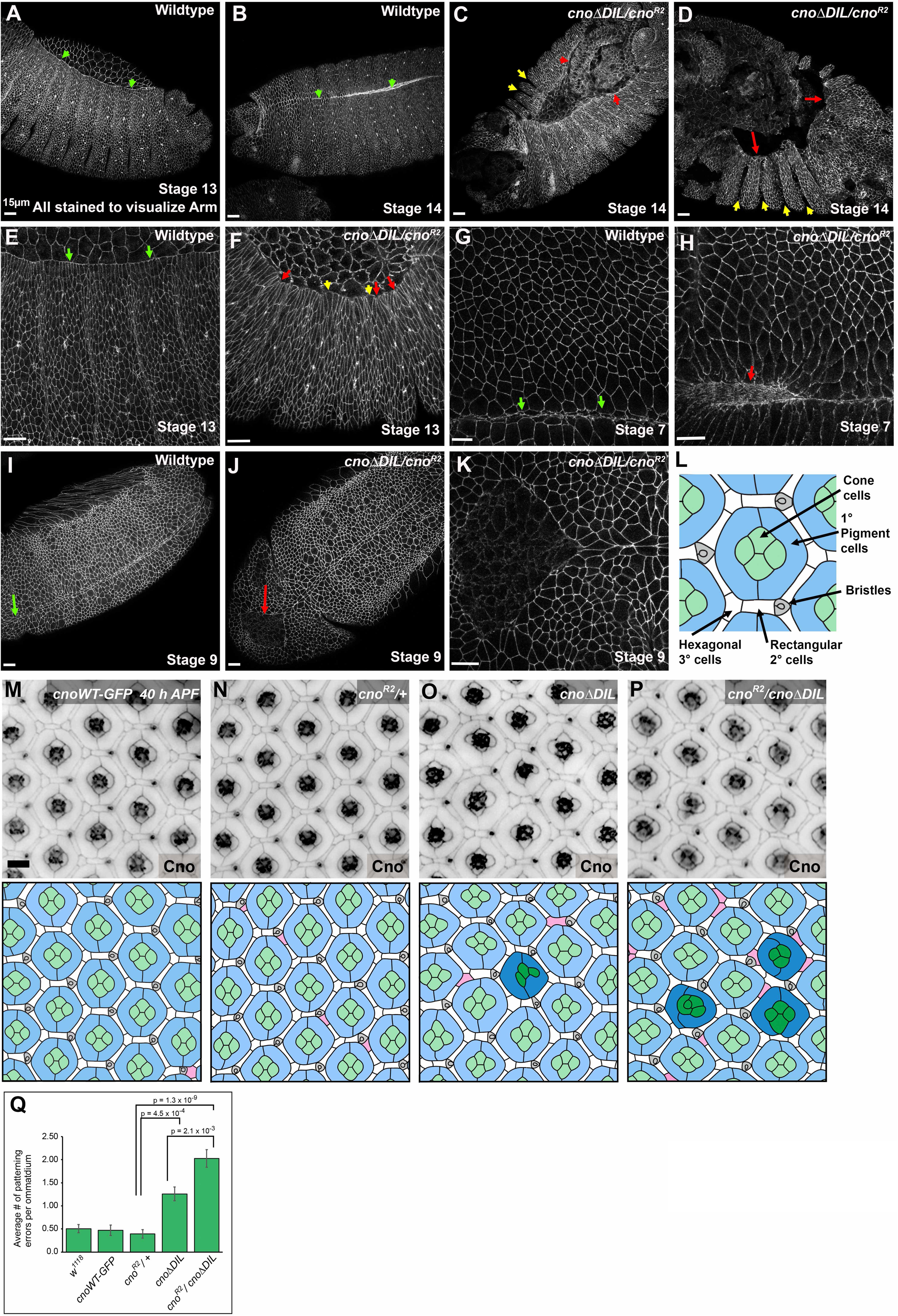
CnoΔDIL does not provide fully wildtype function in embryonic morphogenesis or eye development. (A-K) Embryos, anterior left, stages, and antigens indicated. A-I dorsal up, J,K, ventral view. Embryos labeled *cno*Δ*DIL/cno^R2^*are progeny of the cross of *cno*Δ*DIL/cno^R2^* parents—genotype was not determined. (A-F) Stages 13 and 14 = mid-late dorsal closure. (A, B) During wildtype dorsal closure, as the amnioserosal cells constrict, the lateral epidermis extends dorsally (A, arrows) and zippers closed at the dorsal midline (B, arrows). (C,D) Dorsal closure failure in embryos from the *cno*Δ*DIL/cno^R2^*cross. The epidermal leading edge has detached from the amnioserosa (red arrows), which is undergoing apoptosis before closure is complete. Embryos also exhibit persistent deep segmental grooves (yellow arrows), a characteristic phenotype of strong *cno* mutants. (E) In wildtype leading edge cells extend dorsally, and cell width at the leading edge is relatively uniform (arrows). (F) In embryos from the *cno*Δ*DIL/cno^R2^*cross, some leading edge cells are hyperconstricted (red arrows) and some are splayed open (yellow arrows). (G,H) Stage 7 embryos. In wildtype the mesoderm is fully invaginated at the ventral midline (G, arrows) while in some embryos from the *cno*Δ*DIL/cno^R2^* cross there were mild defects in full mesoderm invagination. (I-K) Stage 9 embryos. In wildtype the ventral furrow fully closed in wildtype (I, arrow)) while some embryos from the *cno*Δ*DIL/cno^R2^* cross had an open ventral furrow anteriorly (J, arrow; K). (L) Cartoon of the pupal eye ommatidium at 40 h APF. (M-P) Small regions of *cno-WT-GFP* (M), *cno^R2^ /+* (N)*, cnoΔDIL-GFP* (O), and *cno^R2^ /cnoΔDIL-GFP* (P) eyes dissected at 40 h APF. GFP-tagged Cno proteins were detected in M, O and P, and endogenous Cno in M. Tracings of each image are presented below, with the cone cells in green, 1° pigment cells in blue, bristle groups in grey, and lattice in white, as indicated in (L). Patterning errors are indicated in darker shades of green or blue, and lattice cells that are incorrectly placed or shaped or in excess in the tissue are highlighted in pink. (Q) Patterning errors, quantified as average number of patterning errors per ommatidium. Scale bar in M=10 µm.

### Cno’s DIL domain is required for Cno’s role in patterning the developing eye

Cell shape change and tissue rearrangement are not confined to embryonic development. The developing eye provides another outstanding place to explore how AJ-cytoskeletal connections shape tissue morphogenesis. The ∼750 ommatidia of the mature eye emerge during pupal development from a neuroepithelium that becomes precisely patterned so that, when examined 40 h after pupal development has begun, each nascent ommatidium has a stereotyped arrangement of epithelial cell types. Each cell type is easily identified due to its characteristic shape (Johnson, 2021). Clusters of four cone cells sit at the center of each ommatidium, surrounded by two primary (1°) pigment cells, and each of the ∼750 ommatidia are separated by a neat lattice of rectangular secondary (2°) and hexagonal tertiary (3°) pigment cells, as well as bristle precursors (Figure 10L, M). Cno is important for the proper development of ommatidial architecture—weak alleles alter the stereotyped arrangement of cells (Gaengel and Mlodzik, 2003; Matsuo et al., 1997; Matsuo et al., 1999), while complete loss of function dramatically disrupts epithelial architecture (Walther et al., 2018).

In our previous work we found that while *cno*Δ*PDZ* and *cno*Δ*FAB* mutants had only subtle defects in embryonic development, they had penetrant defects in cell arrangements in the developing pupal eye (Perez-Vale et al., 2021), suggesting that this tissue provided a more sensitive place to examine protein function. We thus examined pupal eye development in *cno*Δ*DIL* mutants, using wildtype flies (mutant for the *white* gene), flies carrying a GFP-tagged wildtype *cno* gene (Cno-WTGFP), and flies heterozygous mutant for the *cno* null allele *cno^R2^* as controls. *cno*Δ*DIL* homozygous mutant pupae had an elevated number of defective ommatidia with incorrect cell arrangements (Fig. 10O, Q). The cone cells were occasionally mis-configured, ommatidia were sometimes observed with fewer than four cone cells, and the shapes of 1° cells were less ordered (Figure 10O). The lattice was also less precise, with lattice cells occasionally mis-placed or incorrectly shaped. Quantifying total defects, using our previously developed scoring scheme (Johnson and Cagan, 2009), verified the increase in defect frequency (Fig. 10Q)—the defect frequency in *cno*Δ*DIL* homozygotes was roughly comparable to the defect frequency we saw in *cno*Δ*FAB* mutants and somewhat less severe than we observed in *cno*Δ*PDZ* (Perez-Vale et al., 2021). Patterning errors were enhanced in flies in which *cno*Δ*DIL* was heterozygous with *cno^R2^*, thus reducing protein levels (Fig 10P vs 10N, 10O; Fig 10Q), with errors in cone-cell and 1° cell configuration consistent with disrupted cell adhesion during the earlier morphogenesis of these cells. Thus, the DIL domain is important for Cno’s role in eye development.

## Discussion

The dramatic events of embryonic morphogenesis depend on establishing and maintaining robust yet dynamic connections between cell-cell adherens junctions and the actomyosin cytoskeleton. We seek to define the molecular mechanisms by which these connections are made via a network of interconnected proteins. Cno and its mammalian homolog Afadin are central players in this network. In their absence, the complex events of gastrulation and other morphogenetic movement fail, as AJ-cytoskeletal connections are disrupted at places where force is exerted on junctions. Cno provides a superb entry point for defining molecular mechanisms as it, like many AJ proteins, is a complex multidomain protein, with its different domains allowing multivalent connections to diverse other proteins in the network. We are systematically exploring the role of its many folded protein domains in Cno’s biochemical and cell biological functions.

### Canoe’s Dilute domain can dimerize and has a prominent conserved grove on its surface

The DIL domain is the largest of the conserved folded domains in Cno and Afadin, but we had little information about its biochemical or biological functions. The DIL domain in MyoV, the only other protein family in which it occurs, provided speculative possibilities. In MyoV the DIL domain serves two major functions: it acts as an interface for multiple intramolecular and intermolecular interactions, and it can dimerize the protein.

The new tools available from AlphaFold (Jumper et al., 2021; Varadi et al., 2022) provided an opportunity to compare the predicted structure of the DIL domains of Cno and Afadin with that of MyoV. The predicted structures of the Cno and Afadin DIL domains are strikingly similar to those of different MyoV family members, with 15 α-helices forming an elongated domain, as in the MyoV DIL domain. Differences between MyoV and Cno/Afadin are largely confined to a subset of the interhelical loops and the C-terminal 15th alpha-helix. Cno and Afadin are even more similar to one another in predicted structure, with only a few minor differences in interhelical loops. Examining sequence conservation between Drosophila Cno and mammalian Afadin provided additional insights. Intriguingly, the regions that mediate the many intramolecular and intermolecular interactions of MyoV are not well conserved between Cno and Afadin. This includes the region of MyoV that binds Rab GTPases, which was of particular interest because of the known regulation of Cno and Afadin by the small GTPase Rap1. Instead, while Cno and Afadin share only 48% overall identity, there is very strong conservation of a groove on the surface of the DIL domain consisting of residues from α6, α7, α9, and α10 that are distinct from those found in MyoV. We suspect this groove serves as a protein interaction site, and it will be of interest to determine the nature of the potential intramolecular or intermolecular interactions occuring here.

We now have in hand structures or predicted structures of all five folded protein domains that make up the N-terminal half of the Cno/Afadin proteins. What we lack is any information about how they interact with one another. We do know that the N- and C-termini of the DIL domain are in close proximity on one end of the elongated predicted structure, and that the linker between the DIL domain and the PDZ domain that immediately follows is quite short (6-10 amino acids). If the PDZ domain links Cno to the C-terminal tails of transmembrane adhesion receptors like E-cadherin and nectins, that will place the DIL domain relatively close to the plasma membrane. We are keenly interested in using either structural or computational approaches to see if these five domains and their binding partner, Rap1, occur in one or several quaternary conformations that might regulate activity. However, the fact that one can delete either the DIL domain or the PDZ domain without drastically disrupting Cno function calls into question the importance of hypothetical structures such as these, unless they can accommodate a major structural change.

One property conferred on MyoV by its DIL domain is the ability to dimerize. We wondered whether Cno’s DIL domain shared this property. Using Size Exclusion Chromatography and Multi-Angle Light Scattering (SEC-MALS), we found that the purified DIL domain of Cno can dimerize in vitro. We observed both monomer and dimer populations, suggesting a relatively weak interaction. However, in AJs clustering might increase local concentrations, favoring dimerization. This property may help explain the ability of both Afadin and Cno to dimerize/oligomerize in vivo (Bonello et al., 2018; Mandai et al., 1997). However, we suspect interactions in the AJ complex are multivalent, including both direct and indirect interactions among the proteins in the network, and thus Cno and Afadin may self-associate via multiple means.

### Cno recruitment to AJs involves multivalent interactions, as deleting individual protein domains does not prevent it

Our CRISPR-based genetic platform allowed us to replace the *cno* coding sequence at the locus with a version that cleanly lacked the DIL domain. Loss of the DIL domain did not alter recruitment to AJs from their initial establishment to the end of morphogenesis. It also did not alter enrichment of Cno protein at the AJs which are under elevated tension. In this way, it resembled two of the mutants we examined earlier, *cno*Δ*PDZ* and *cno*Δ*FAB* (Perez-Vale et al., 2021). While the tandem RA domain cassette and its Rap1 binding partner are important for initial Cno localization to AJs as they first assemble, they become dispensable for AJ localization after gastrulation onset (Perez-Vale et al., 2023; Perez-Vale et al., 2021). Thus, four of the five folded domains present in Cno, along with its conserved C-terminal FAB, are each singly non-essential for AJ recruitment, despite their conservation from Drosophila to mammals, over ∼600 million years of evolutionary time. These data reinforce the remarkable multivalent nature of AJ assembly, with multiple interactions appearing to be sufficient for Cno recruitment to junctions. In future studies, it will be important to test the FHA domain and the remainder of the intrinsically disordered region—perhaps one of those might be essential for AJ localization. Moving on, mutating multiple domains simultaneously might help reveal potential redundancy.

### Canoe’s DIL domain is not essential for viability but plays a supporting role in Cno function in morphogenesis

The conservation of the DIL domain in all animal family members led us to suspect it would play an important role in Cno/Afadin function in vivo. However, to our surprise, *cno*Δ*DIL* mutants are viable and exhibit normal or near normal fertility. Closer examination of *cno*ΔDIL mutants during morphogenesis verified normal function during cellularization, germband elongation and dorsal closure, times at which AJs need to dynamically respond to tension as cells change shape. It was only when we reduced CnoΔDIL levels, using our sensitized assay in which we examined the progeny of *cno*Δ*DIL/cno^R2^* parents, that its importance in maintaining robustness became apparent. CnoΔDIL could not provide full function in embryos lacking maternal and zygotic wildtype Cno, and thus the morphogenesis defects of *cno* zygotic null mutants were substantially enhanced, with increased failure of dorsal closure, and failure to fully internalize the mesoderm.

How can a protein domain that has been maintained through 600 million years of evolutionary divergence (Gurley et al., 2023) be dispensable for embryonic morphogenesis? Together with our earlier work on *cno*Δ*PDZ* and *cno*Δ*FAB* (Perez-Vale et al., 2021), these data emphasize the power of natural selection to maintain protein structure even when mutants appear wildtype or near wildtype in our lab-based assays. In the case of *cno*Δ*FAB* mutants, the 25% embryonic lethality would provide ample scope for selection. We think it is likely that *cno*Δ*PDZ* and *cno*Δ*DIL* mutants have defects in viability and morphogenesis that are within the noise of our phenotypic analysis, and that the activity that these domains confer was sufficient to maintain Cno/Afadin protein structure and to maintain substantial sequence conservation. Further, *cno*Δ*DIL* mutants share defects in the precision of eye development with cnoΔ*PDZ* and *cno*Δ*FAB* – these and potential similar issues like this in other internal and/or postembryonic tissues are also likely to be sufficient to empower selection. We imagine similar constraints explain the maintenance of genes like *sidekick* and *vinculin,* in which animals completely lacking these proteins are viable and fertile—*sidekick* mutants have defects in ommatidial development similar to those we observed here (Letizia et al., 2019). Together, these emphasize the need for robustness in AJ-cytoskeletal connections during multiple embryonic and postembryonic events, and the need to experimentally map and characterize the multivalent interactions that underlie the network.

## Author Contributions

Emily McParland carried out genetic and cell biological analyses of the *cno*Δ*DIL* mutant, T. Amber Butcher designed the *cno*Δ*DIL* mutant and carried out biochemical analysis of the DIL domain with advice from Kevin Slep, Noah Gurley carried out experiments to validate the allele and analyzed levels of protein expression, Kevin Slep used AlphaFold data to examine conservation of the DIL domain, Ruth Johnson analyzed phenotypes in the developing eye, Mark Peifer assisted with Drosophila genetics, and Emily McParland, T. Amber Butcher, Noah Gurley, Ruth Johnson, Kevin Slep and Mark Peifer wrote the manuscript.

## Acknowledgements

We are grateful to Dr. Ashutosh Tripathy for assistance with SECMALS, to Jenevieve Norton for help with the sensitized assay, and to Peifer, Bergstralh, Finegan, and Williams lab members for helpful discussions. Work in the Johnson lab is supported by R15 GM114729 to R.I. Johnson. This work was funded by NIH R35 GM118096 to M. Peifer.

## Materials and Methods

### Cloning and Purification of Cno DIL Domain Construct

The Cno DIL domain sequence (aa 613-1006) was cloned into plasmid pET28 (Millipore Sigma, Burlington, MA) using PCR and primers with engineered NheI and EcoRI restriction sites. Plasmid containing the construct was transformed into BL21 (DE3) pLysS E. coli cells and grown in 8 liters LB media with 20 mg/l kanamycin at 37°C. After reaching an optical density of 0.8 at 600 nm protein expression was induced with 100 μM IPTG for 16 hours at 20°C. Cells were harvested by centrifugation at 4°C, then were resuspended in 200 ml buffer A (25 mM Tris (pH 8.5), 300 mM NaCl, 0.1% β-mercaptoethanol, 10 mM imidazole) supplemented with 5 µg/mL DNase, 10 µg/mL lysozyme, and 0.5 mM PMSF. Cells were lysed by sonication. Lysate was clarified by centrifugation at 23,000 x g for 45 minutes at 4°C. Supernatant was loaded onto a nickel-nitriloacetic acid (Ni^2+^-NTA) column (Qiagen, Hilden, Germany), washed with 500ml buffer A, and eluted in 75 ml buffer B (buffer A + 290 mM imidazole) to which 25 μl thrombin (6 mg/ml)(Prolytix, Essex Junction, VT) and CaCl_2_ (1 mM final concentration) was added. Cleavage of the His_6_ tag with thrombin digestion occurred overnight at 4°C. Cno DIL domain was dialyzed overnight against 25 mM Tris (pH 8.5), 300 mM NaCl, and 0.1% β-mercaptoethanol, filtered successively over benzamidine Sepharose (Cytiva, Marlborough, MA) and Ni^2+^-NTA resin. Protein was exchanged into 25 mM Tris (pH 8.5), 300 mM NaCl, and 0.1% β-mercaptoethanol, concentrated in a 30,000 MWCO Millipore concentrator to 85.2 mg/ml, and flash frozen in liquid nitrogen for storage at −80°C.

### SECMALS Data Collection

Purified and concentrated Cno DIL domain protein was thawed at room temperature and diluted to 5.0 mg/ml in 300mM NaCl, 25mM Tris (pH 8.5), 1mM MgCl_2_, 0.1% β-mercaptoethanol, and 0.2 g/l sodium azide to a total volume of 120 μl. 100 μl of diluted protein was injected onto a Superdex 200 column with a flow rate of 0.5 ml/min. The protein passed through a Wyatt Optilab rES refractometer followed by a DAWN HELEOS II light-scattering instrument. Refractive index and light-scattering data were processed using the Astra software program and were used to determine the molar mass of the protein. The SEC-MALS data shown is representative of experiments conducted in duplicate using a biological replicate.

### Structure Prediction and Structure Analysis

Drosophila Cno and rat Afadin structure prediction models were obtained from the AlphaFold server (Jumper et al., 2021; Varadi et al., 2022). The MyoVb DIL domain was obtained from PDB 4J5M (Nascimento et al., 2013). Coordinates of MyoV binding proteins in complex with the MyoV DIL domain are from PDB 4KP3 (RILPL2 (Wei et al., 2013), 4LX0 (Rab11 (Pylypenko et al., 2013)), 4KP3 (Melanophilin (Wei et al., 2013), 5JCY (Spir2 (Pylypenko et al., 2016)), and 6KU0 (MICAL1 (Niu et al., 2020)). Coordinates of the full length MyoVa structure are from PDB 7YV9 (Niu et al., 2022)). Structures were aligned using the align command in PyMOL (Schrodinger). Protein sequence alignments were generated using Clustal Omega (Madeira et al., 2022), and adjusted manually based on output from the PyMOL structural alignments.

### Fly work

We used *yellow white* flies as our control and we refer to them in the text as wild type. All the experiments were performed at 25°C.

### Generating the mutant rescue construct and its ΦC31-mediated integration into the *cno*ΔΔ allele attP site

The *cnoWT-GFP* rescue construct (Perez-Vale et al., 2021) was used to generate a new construct lacking the DIL domain. To remove this domain we designed an in-frame deletion of amino acids 613-993 of *Drosophila* Cno. We created this deletion using a Q5 Site-Directed Mutagenesis kit (New England Biolabs, Cat. No: E0554S) with subsequent sequence verification. The vector carrying the *cno*Δ*DIL* gene, pGE-attB-GMR, also carries a w^+^ selectable marker next to the *cno* coding sequence, and both are flanked by attR and attL sites allowing site-specific integration into the attP site at the *cno*ΔΔ locus (Perez-Vale et al., 2021). Injection of the *cno*ΔDIL-GFP rescue construct was carried out by BestGene (Chino Hills, CA)—this DNA was injected into *PhiC31/int^DM.Vas^; cno*ΔΔ embryos. F1 offspring were screened for the presence of the w^+^ marker and outcrossed to *w; TM6B, Tb/TM3, Sb* to generate a balanced stock over TM3. We verified the integration of *cno*Δ*DIL-GFP* by both PCR amplification and sequencing and by immunoblotting. To remove potential other mutations from the *cno*Δ*DIL-GFP* chromosome we outcrossed the stock to a *y w* stock with a wildtype 3rd chromosome for multiple generations, selecting for the linked w^+^ marker in each generation. This allowed us to homozygose *cno*Δ*DIL-GFP*.

### Molecular characterization of engineered *cno* allele

The following primers were used for PCR amplification of the sequence between the intron upstream of construct insertion and the exon after RA1: forward (F.1), 5’-ACCGTCACAAACCAACCAGA-3’; reverse (R.1), 5’-AACACCATTTCCCAAGCCCA-3’. The RA1 domain is encoded by two exons which, in the wildtype CDS, are separated by an intron sequence. Because ΔDIL mutants lack wildtype introns after the 5’ UTR, the amplicon produced from ΔDIL mutants is expected to be smaller in size compared to that of wildtype. For PCR amplification of the sequence between the end of FHA and the first PDZ exon, the following primers were used: forward (F.2), 5’-GTGGTAATGTTCGGACGGGT-3’; reverse (R.2), 5’-CCAGGCACCACACTCTTGAT-3’. In the wildtype CDS, there exist several introns between the FHA and the beginning of the PDZ domain. Again, because ΔDIL mutants lack introns after the 5’ UTR, the amplicon produced from ΔDIL mutants is expected be smaller in size compared to that of wildtype. Finally, for PCR amplification of the sequence between the end of FHA and the intron after the first DIL exon, the following primers were used: forward (F.3), 5’-GTGGTAATGTTCGGACGGGT-3’; reverse (R.3), 5’-AATGGCGGCTGCTTTCAT-3’. In this pairing, since the reverse primer targets an intron sequence that is non-existent in ΔDIL mutants, an amplicon is only produced for wildtype.

### Embryo Fixation and Immunofluorescence

Eggs were collected in cups at 25°C on apple juice agar plates with yeast paste. Embryos were dechorionated in 50% bleach, washed three times in 0.03% Triton X-100 with 68 mM NaCl, and then fixed in 95°C Triton salt solution (0.03% Triton X-100 with 68 mM NaCl, 8 mM EGTA) for 10 seconds. We then added ice-cold triton salt solution and transfer to ice for fast cooling for at least 30 minutes. We devitellinized the embryos by vigorous shaking in 1:1 heptane:methanol solution. The embryos were then washed three times with 95% methanol/5% EGTA. After this step embryos were sometimes stored in 95% methanol/5% EGTA at −20°C overnight before staining. Before staining, the embryos were washed three times with 5% normal goat serum/0.1% saponin in phosphate-buffered saline (PBS) (PBSS-NGS). We then blocked in 1% normal goat serum (NGS) in PBSS-NGS for 1 hour, and embryos were incubated in primary antibodies overnight at 4°C or 2-3 hours at room temperature. Once incubation finished, we washed three times with PBSS-NGS and incubated embryos in secondary antibodies overnight at 4°C or 2-3 hours at room temperature. Both primary and secondary antibodies were diluted in 1% bovine serum albumin/0.1% saponin in PBS, and the dilutions used are listed in Table 3. After the secondary antibody incubation, we washed three times with PBSS-NGS and stored embryos in 50% glycerol until mounted on glass slides using a homemade Gelvatol solution (recipe from the University of Pittsburg’s Center for Biological Imaging). Antibodies used are in Table 2

**Table 1.**

|  |  | Expected product size |  |
| --- | --- | --- | --- |
| *Primer pair* |  | Edited locus | Wild-type locus |
| 1 | F.1 (Forward.1) | 1346 bp | 1680 bp |
|  | R.1 (Reverse.1) |  |  |
| 2 | F.2 (Forward.2) | 588 bp | 4238 bp |
|  | R.2 (Reverse.2) |  |  |
| 3 | F.3 (Forward.3) | No product | 1769 bp |
|  | R.3 (Reverse.3) |  |  |

**Table 2.** Antibodies used in this study.

| Primaries | Species | Dilution | Source |
| --- | --- | --- | --- |
| Anti-Canoe | Rabbit IgG | 1:1,000 (IF, WB) | Sawyer et al., 2009 |
| Anti-Bazooka | Rabbit IgG | 1:2,000 (IF) | Choi et al., 2013 |
| Anti-Armadillo (N27A1) | Mouse IgG<sub>2a</sub> | 1:100 (IF) | Developmental Studies Hybridoma Bank |
| Anti-GFP (JL-8) | Mouse IgG<sub>2a</sub> | 1:1,000 (IF, WB) | Clontech Laboratories (632381) |
| Anti-α-tubulin | Mouse IgG<sub>1</sub> | 1:5,000 (WB) | Sigma-Aldrich (T6199) |
| **Secondary antibodies** |  | **Dilution** | **Source** |
| Alexa Fluor 488, 568, 647 |  | 1:1,000 (IF) | Life Technologies |
| Anti-Rabbit IRDye 680RD |  | 1:10,000 (WB) | LI-COR Biosciences |
| Anti-Mouse IRDye 800CW |  | 1:10,000 (WB) | LI-COR Biosciences |

IF, immunofluorescence; WB, Western blot.

### Image Acquisition and Analysis

Fixed embryos were imaged on a confocal laser-scanning microscope (LSM 880; 40x/NA 1.3 Plan-Apochromat oil objective; Carl Zeiss, Jena Germany). Images were processed and maximum intensity projections were generated using ZEN 2009 software. We used Photoshop (Adobe, San Jose, CA) to adjust input levels and brightness and contrast. Analysis of apical-basal positioning on maximum intensity projections (MIPs) was executed as previously described (Choi et al., 2013). Briefly, using Zen 2009 software, the z-stacks were cropped to select a region of interest (ROI) on the *xy*-axis of 250×250 pixels for stacks collected using a digital zoom of 2 or 200×200 pixels for stacks with a 1.6 digital zoom. Using the Zen software the z-stacks ROIs dimensions were modified from *yzx* to *xyz* along the *y*-axis from which MIP were generated.

### Cno SAJ and TCJ enrichment analysis

Data for analyzing SAJ and TCJ enrichment was obtained from z-stacks taken through the embryo using a digital zoom of 1.6 or 2 and a step size of 0.3 µm. First, the total length of the cells was defined by determining slice position of the apical (top) and basal (bottom) of the cells in the stack. For embryos in mid-late stage 5 the SAJs were determined to be at 21.82% of the total length and the TCJ enrichment was assessed at 33.33% more basal to the SAJs. For embryos in late stage 5, the SAJs correspond to 21.82% of the total length and the TCJ enrichment was assessed at 50% more basal to the SAJs.

The Cno TCJ intensity ratio was measured from MIPs of a 1.2-2.4 µm of the apical AJ region of embryos from stage 7. The MIPs were generated from z-stacks taken through the embryo using a digital zoom of 1.6 or 2 and a step size of 0.3 µm. ImageJ software was used to identify the apical AJ region from which z-stack MIPs were generated. The mean intensity of Cno was measured using FIJI (National Institutes of Health, Bethesda, MD, USA) by creating lines using the line tool (line width of 5 pixels) along the bicellular junctions, avoiding TCJs or multicellular junctions, and next creating short lines at TCJ/multicellular junctions at 300% zoom. For each TCJ or four-way junction, the three or four bicellular junctions in contact with that junction were measured to obtain the mean intensity. For each junction, a short line was drawn in the cytoplasm to standardize pixel intensity for the image by subtracting cytoplasmic background from the junctional intensity. A total of ten cells were quantified per embryo and a total of four embryos were assessed from three experiments. The average bicellular junction intensity per cell was calculated. The Cno TCJ ratio was calculated by dividing the mean intensity of the TCJ by the average of the bicellular junctions. Box and whiskers graphs were made using GraphPad: the box shows the 25^th^-75^th^ percentile, the whiskers are 5^th^-95^th^ percentiles, the horizontal line is the median and the plus sign (+) is the mean. Data statistical analysis was done using GraphPad. Statistical significance was calculated by Welch’s unpaired t-test or Brown Forsythe and Welch ANOVA test.

### Planar polarity quantification

The planar polarity of Cno was measured from MIP of a 2.4 µm region of the apical AJs of embryos from stage 7 to early stage 8. The MIPs were generated from z-stacks taken through the embryo using a digital zoom of 1.6 or 2 and a step size of 0.3 µm. Using FIJI the 2.4 µm region was identified from which MIPs were generated. The mean intensity was measured using FIJI by creating lines (line width of 5 pixels) at 300% zoom at bicellular borders, without including the TCJ/multicellular junctions. Anteroposterior and dorsoventral borders were selected manually, choosing cells with aligned anterior-posterior borders where planar polarity is most apparent. The background intensity was measured by drawing a short line in the cytoplasm of all cells measured. The background pixel intensity was subtracted from AP and DV border intensities. A total of four embryos were assessed from at least four experiments. Cno was normalized to DV borders producing an AP/DV ratio. Box and whiskers graphs were made using GraphPad. The box shows the 25^th^-75^th^ percentile, the whiskers show 5^th^-95^th^ percentiles, the horizontal line shows the median and the plus sign (+) is the mean. Data statistical analysis was done using GraphPad. Statistical significance was calculated using a one-way ANOVA.

### Cuticle preparation and analysis

We prepared embryonic cuticles according to (Wieschaus and Nüsslein-Volhard, 1986). Embryos were collected on apple juice agar plates with yeast, aligned on a fresh apple juice agar plate without yeast and incubated at 25°C for 48 hours to allow embryos to develop fully and viable embryos to hatch. All unhatched embryos were collected in 0.1% Triton X-100 and dechorionated in 50% bleach for 5 minutes. They were then washed three times with 0.1% Triton X-100 and transferred to glass slides, where all the liquid was removed, mounted in 1:1 Hoyer’s medium:lactic acid, and incubated at 60°C for 24-48 hours. They were then stored at room temperature. Images were taken using a Nikon Labophot with a 10x Phase 2 lens, and captured on an iPhone, and placed into categories based on morphological criteria.

### Western blotting

Table 2 contains the antibodies and dilutions used for these experiments. Protein levels expression of Cno, Pyd, and Arm were determined by immunoblotting embryos collected in the 1-4 hours and 12-15 hours windows. The lysates were generated as in (Manning et al., 2019). Briefly, embryos were dechorionated for 5 minutes in 50% bleach. After washing three times with 0.1% Triton X-100, lysis buffer (1% NP-40, 0.5% Na deoxycholate, 0.1% SDS, 50 mM Tris pH 8.0, 300 mM NaCl, 1.0 mM DTT, 1x Halt protease, phosphatase inhibitor cocktail (100x), and 1 mM EDTA) was added and the embryos were placed on ice. Embryos were ground in a microcentrifuge tube using a pestle, lysate was centrifugated at 13200 RPM for 15 minutes at 4°C, and protein concentration was determined using Bio-Rad Protein Assay Dye. The lysates were resolved using 7% SDS-PAGE and transferred onto nitrocellulose membranes with a pore size of 0.2 μm. The membranes were blocked in 10% bovine serum albumin (BSA) diluted in tris-buffered saline with 0.1% Tween-20 (TBST) for 1 hour at room temperature. For primary and secondary staining, antibodies were diluted in 5% BSA with TBST. Incubation was performed either for 2 hours at room temperature or overnight at 4°C for the primary antibody, and a 45-minute incubation at room temperature was performed for the secondary antibody. The membranes were developed using the Odyssey CLx infrared system (LI-COR Biosciences). Analysis of band densitometry was calculated using Empiria Studio® Software.

### Pupal eye dissection, immunofluorescence and analysis

Wildtype and mutant stocks were maintained on nutrient-rich Drosophila media at 25°C. Pre-pupae were selected and maintained in humidified chambers until dissection at 40 hours after puparium formation (h APF) (DeAngelis and Johnson, 2019). Rabbit anti-Cno (1:500), and chicken anti-GFP (1:8000, Abcam #13970) followed by were Alexa 488-conjugated secondary antibodies (Jackson ImmunoResearch #711-545-152 or #703-545-155) were used to detect Cno, Cno-WT-GFP and *cno*Δ*DIL-GFP*, and retinas imaged with a Leica DM5500 B fluorescence microscope. We performed dissections in triplicate, with 5-10 pupae of each genotype dissected each time. Patterning errors were scored in retinas from one representative triplicate, as previously described (Johnson and Cagan, 2009). Analyses spanned 9-15 eyes for each genotype, with 110 data points per genotype. Image files were processed for publication using Adobe Photoshop.

